# Zoonotic infections and genomic evolution of novel reassortant swine influenza A viruses in Spain

**DOI:** 10.64898/2026.05.22.724525

**Authors:** Paloma A. Encinas, Brady O’Boyle, Alexander Maksaiev, Martha I. Nelson, Adolfo García-Sastre, Gustavo del Real

**Author notes:** Corresponding Authors: Paloma A. Encinas, Martha I. Nelson.

## Abstract

Influenza A virus (IAV) circulates widely in European pig populations and continues to diversify through frequent introductions from humans, followed by reassortment within swine. Spain represents a particularly dynamic ecological setting due to the coexistence of intensive white⍰pig production, extensive Iberian⍰pig systems, and abundant wild boar populations. This study provides an integrated analysis of IAV evolution and genomic diversity in swine in Spain between 2019 and 2022, expanding on previous surveillance from 2016 to 2019. Sampling across 24 provinces yielded 66 new whole⍰genome sequences from Iberian and white pigs.

We identified 18 genotypes, including 11 novel reassortants not detected in our previous survey. Several genotypes, such as H1huN2 G21 and G22, H3N2 G23, and the unusual H3N1 G12, were exclusive to the country. Some genotypes were detected across white pigs, Iberian pigs, and wild boar in Toledo and Badajoz, suggesting viral flow among swine populations.

Phylogenetic analyses revealed ongoing introductions of H1N1pdm09 from humans into pigs, generating at least five reassortant genotypes (G10, G16–G19). These lineages incorporated pandemic internal cassettes and, in some cases, human⍰seasonal N2 segments, highlighting the continued role of humans as a source of viral incursions. Conversely, four zoonotic infections (H1N1v) detected in Spain between 2022 and 2026 were linked to genotypes circulating in white pigs, underscoring the bidirectional nature of IAV transmission at the human swine interface.

Overall, this study demonstrates that Spain provides ecological conditions conducive to IAV diversification, reassortment, and zoonotic risk. The findings reinforce the need for sustained One Health surveillance.

**Highlights:**

- Novel swine influenza virus (SIV) genotypes exclusive to Spain
- Phylogenetic analysis of genomic segments of zoonotic variants of swine origin detected in Spain since 2022
- Shared circulation of influenza A compatible with interbreed transmission among domestic pigs and wild boar

## 1. Introduction

Swine influenza is caused by the influenza A virus (IAV), a member of the Orthomyxoviridae family, characterized by a segmented RNA genome. This virus primarily causes respiratory disease in pigs, but is also capable of infecting other vertebrates such as humans and birds (1). In addition to the circulating swine IAV, pigs can be infected with IAV from other host species, as they possess receptors for both human and avian⍰origin IAVs in their respiratory tract (2). When IAVs from different origins, especially swine, avian and human, co-infect a pig, they can exchange genetic segments and produce reassortant viruses, which can establish new viral genotypes lineages that may have greater virulence in the swine population (3).

IAVs can be classified into subtypes, based on the main surface proteins of the virus: haemagglutinin (HA) and neuraminidase (NA). The most frequently detected swine influenza virus (SIV) subtypes are: H1N1, H1N2, H3N2 and H1N1pdm09 (4, 5). IAVs can be further classified based on the evolutionary history of the segments. Regarding IAV lineages in Europe, three distinct haemagglutinin H1 lineages are currently circulating: the Eurasian avian-like lineage (EAswH1, H1av), the human seasonal-like lineage (HUswH1, H1hu), and the pandemic-derived lineage (PDMswH1, H1p) (6, 7). In addition, there are two human seasonal-like H3 lineages circulating in Europe: an H3 lineage from 1970s (1970s-like H3) and a new H3 lineage that emerged in the 2000s (2000s-like H3) (8, 9). Epidemiological surveillance is the cornerstone for early detection of emerging, new, more virulent IAVs (10). In spite of its relevance, coordinated studies on SIV surveillance in the European swine population were discontinued after 2013 (11). Since then, studies have been conducted in different European countries revealing a high genetic variability of SIVs, with regional differences in the frequency of HA subtypes and lineages (12–18).

Spain is among the world’s leading producers of commercial white pigs and is the main source of Iberian pigs, an autochthonous breed of the Iberian Peninsula which accounts for 10% of the countrýs total pork production (19, 20). White pigs are typically raised under intensive production systems, where IAV is enzootic and the probability of human-to-pig transmission is high due to frequent contact with farm workers (21–24). In contrast, Iberian pigs are raised in extensive systems, roaming freely in the Dehesas, particularly concentrated in Extremadura, which maintains 38% of national production (25), and consuming natural forage and acorns during the montanera season (October–March), where they interact with wildlife, including birds and wild boars, thereby increasing exposure to other pathogens, including IAVs, of heterogeneous origin (26, 27). This ecological context positions Spain as a potential key setting for IAV diversification, where viruses from multiple host reservoirs can converge in pigs, facilitating reassortment and the emergence of novel strains.

Frequent human-to-pig transmissions have been reported worldwide (21, 28–31) and, in recent years, pig-to-human transmissions, known as human influenza variants (v), have taken place in the European region (31–35), four of them in Spain (35, 36) since 2022. The independent circulation of previous human IAVs in swine populations has been classified as a significant risk of zoonotic reintroduction of antigenically modified IAVs into the human population, which may evade human immunity and spread more efficiently, potentially leading to a new influenza pandemic like the one in 2009 (37).

We previously characterized the genetic diversity of swine influenza viruses (SIV) circulating in Spain between 2016 and 2019 (38), identifying twelve distinct genotypes encompassing five haemagglutinin lineages and four subtypes. To conduct robust risk assessments of influenza infections in pigs, from both an animal and public health standpoint, it is essential to continue monitoring the evolution of SIV, given its continuously increasing genetic and antigenic complexity.

Our objective is to investigate the evolution of SIVs circulating in Iberian and white pigs in Spain during the 2019–2022 period, with particular emphasis on the emergence of newly introduced subtypes at the regional level and the detection of novel reassortant viruses. Additionally, we aim to further elucidate the phylogenetic origin of the human variant cases reported in Spain and assess their relationship to the SIVs currently circulating in the country.

## 2. Materials and methods

### 2.1. Sample collection

Sampling was opportunistic, relying on animals submitted for routine diagnostic investigation rather than on a structured or pre-planned sampling design. Samples were obtained via passive surveillance or through the collection of respiratory tissues during diagnostic necropsies.

Passive sampling in Iberian pigs consisted of respiratory specimens and sera collected by farm veterinarians in Extremadura, the main Iberian pig⍰producing region in Spain. A total of 288 nasal swabs and 603 serum samples were obtained from 25 Iberian pig farms during 2020. Additional samples, consisting of bronchial scrapings and post-mortem lung tissues that tested PCR positive for influenza A, were retrieved from Iberian and white pigs submitted for necropsy as part of routine veterinary diagnostics between 2019 and 2022, in order to increase the number of SIV isolates detected in Spain. In total, 108 samples from 24 provinces (Albacete, Ávila, Badajoz, Barcelona, Burgos, Castellón, Cuenca, Gerona, Huesca, Lérida, Madrid, Málaga, Murcia, Navarra, Palencia, Pontevedra, Salamanca, Segovia, Sevilla, Teruel, Toledo, Valencia, Zamora, and Zaragoza) were collected.

#### Ethics approval statement

No specific ethical approval was required for this study, in accordance with Directive 2010/63/EU of the European Parliament on the protection of animals used for scientific purposes, as all samples were obtained from non-experimental clinical veterinary procedures. No animals were manipulated or euthanized specifically for the purposes of this study.

### 2.2 Serological analysis

Iberian pig serum samples were screened for antibodies against IAV using a commercial indirect enzyme-linked immunosorbent assay (ELISA) targeting the viral nucleoprotein (NP), according to the manufacturer’s instructions (INGEZIM Influenza A, Eurofins-Ingenasa, Spain).

### 2.3 RNA extraction and SIVs Isolation

Nasal swabs were collected in viral transport medium (VTM), transported refrigerated to the laboratory, briefly vortexed, and centrifuged. Lung samples were kept frozen until processing, then homogenized in Phosphate-Buffered Saline (PBS) using a tissue grinder (ULTRA-TURRAX Tube Drive, IKA, Germany) and centrifuged. The resulting supernatants were used for RNA extraction and for inoculation of cell cultures and embryonated chicken eggs for virus isolation.

Total RNA from samples was extracted using the QIAamp Viral RNA Kit in a QIAcube workstation (Qiagen). IAV detection was made by real-time reverse transcription quantitative polymerase chain reaction (RT-qPCR) of Matrix (MP) gene with the primers forward: GAC CRA TCC TGT CAC CTC TGA C, reverse: AGG GCA TTY TGG ACA AAK CGT CTA, and probe: TGC AGT CCT CGC TCA CTG GGC ACG (labeled with 6-FAM, MGBNFQ; Invitrogen). The reaction conditions were in accordance with the World Health Organization (WHO) Manual for the laboratory diagnosis and virological surveillance of influenza (39): RT (45°C, 30□min), activation Taq (95°C, 10□min) followed by 40 cycles: denaturation (95°C, 15□sec) and annealing/extension (60°C, 60□sec). The RT-qPCR was performed with the High ScripTools-Quantimix Easy Probes kit (Biotools) in an Applied Biosystems TM Fast 7500 thermal cycler. IAV isolation from RT-PCR-positive samples was made in Madin–Darby canine kidney cells cultures grown in advanced Dulbecco’s Modified Eagle Medium (DMEM) (Thermo Fisher) supplemented with glutamine (4□µM), 4-(2-hydroxyethyl)-1-piperazineethanesulfonic acid (HEPES) (20□µM), Bovine Serum Albumine (BSA) fraction V, 100□U/ml streptomycin, 100□U/ml penicillin and (6-(1-tosylamido-2-phenyl)-ethyl-chloromethyl-ketone)-treated (TPCK) trypsin (2□µg/ml) from bovine pancreas (Merck Life Science) at 37°C and 5□per cent CO2 according to the Manual on Animal Influenza Diagnosis and Surveillance. Alternatively, if isolation in cell culture was unsuccessful, RT-PCR-positive samples were inoculated in 10-day-old chicken embryos. Before inoculation in embryos and cells, swab fluids and lung extracts in VTM were spinning 4000□rpm, 10□min, and filtered through 0.22-mm Millex syringe filters (Millipore). Inoculated cell cultures were cultured at 37°C, 5□per cent CO2, and examined daily for evidence of cytopathic effects (CPEs). If CPE is positive, or after 3□days, supernatants were tested by RT-PCR. Allantoic fluid was tested for the presence of virus by hemagglutination assay or RT-PCR. Negative allantoic fluid and cell supernatants were passaged up to three times. Virus isolates were submitted to genetic analyses to determine the subtype by conventional one-step RT-PCR with Superscript III one-step RT-PCR Kit with Platinum Taq polymerase (ThermoFisher). Subtypes were further classified regarding its phylogenetic origin in: avian (av), seasonal human (hu) or pandemic (p) as follows: H1avN1av, H1avN2, H1avN1p, H1pN1p, H1pN2, H3N2, H1huN1, H1huN2, H3N1, H3N2.

### 2.4 Whole genome sequencing

Whole genome sequencing of the SIV isolates was performed at the Icahn School of Medicine at Mount Sinai of New York (ISMMS). RNA was extracted using a QIAamp viral RNA kit (Qiagen). The entire genome was amplified from 5□µl of RNA template using a multisegment RT-PCR strategy (40). Multiplex-RT-PCR amplification was performed with a SuperScript III high-fidelity RT-PCR kit (ThermoFisher) according to the manufacturer’s instructions using the Opti primer mix, consisting of primers Opti1-F1 (5’-GTTACGCGCCAGCAAAAGCAGG), Opti1-F2 (5’-GTTACGCGCCAGCGAAAGCAGG), and Opti1-R1 (5’-GTTACGCGCCAGTAGAAACAAGG). DNA amplicons were purified using an Agencourt AMPure XP 5□ml kit (Beckman Coulter). Sequencing libraries were prepared, and sequencing was performed on a MiSeq instrument (Illumina, Cambridge, UK) with 2 × 150-base paired end reads. Handling of the data for the raw sequence reads and extraction of consensus sequences were performed at ISMMS as described previously (41). All sequences were deposited in the National Center for Biotechnology Information’s (NCBI) GenBank, (Table S1).

### 2.5 Phylogenetic analysis

Phylogenetic trees were inferred for each dataset using the maximum likelihood approach available in IQ-Tree v2.1.4 (42). The substitution model used was General Time Reversible with FreeRate heterogeneity across sites (GTR+R). Branch support was assessed using 1000 replicate ultrafast bootstrap approximation. Outlier sequences based on their temporal signal within the background dataset were identified using TempEst v1.5.3 (43) and removed from the dataset. Tree files were visualized in FigTree v1.4 (44).

Our swine dataset included 66 IAV whole genome sequences (WGS) obtained from Iberian and commercial white pigs for this study during 2019-2022 in Spain, 47 IAV WGS obtained from these populations for a previous study 2016-2019 in Spain, 1 IAV WGS isolated from wild boar (45), and 3219 additional WGS from swine and humans in Europe that were obtained from GISAID on March 25, 2026. 201,289 additional sequences from humans in Europe were downloaded from GISAID on April 8, 2026 for pdm09 trees, H1 human seasonal tree, H3 2000s-like tree and N2 tree. Human sequences for the pdm09 dataset were subsampled to 100 sequences per year between 2009-2025. H1 human seasonal, N2, and H3 2000s-like datasets were subsampled to 50 sequences per year between 1977-2008 for H1 and 1968-2025 for N2 and H3. For the 2009-2010 pdm09 tree, the human background dataset was subsampled to 1250 sequences per year between 2009-2010. The HA lineages of IAV sequences were classified using the Bacterial and Viral Bioinformatics Resource Center (BV-BRC) subspecies classification tool (46) and validated phylogenetically. All other segment lineages were classified phylogenetically. Alignments were generated for the coding regions of each of the eight segments of the IAV genome and separately for the HA and NA subtypes, resulting in ten datasets: PB2, PB1, PA, H1, H3, NP, N1, N2, MP, and NS. MAFFT v7.505 was used with 1000 refinement iterations (47). Sequences were trimmed to the start and stop of coding regions, except for the MP and NS segments, which were trimmed from the start of the first coding region to the stop of the second coding region.

Data are available at: https://github.com/boboyle/Zoonotic-infections-in-Spain-associated-with-novel-reassortant-swine-origin-influenza-A-viruses

## 3. Results

### 3.1 Seroprevalence and characterization of SIV from Iberian pigs during the period 2019-2022

Eleven out of 25 (44%) Iberian pig farms in Extremadura tested positive for IAV by ELISA, with an IAV seroprevalence of 20.51% of the tested animals (24/117) in Cáceres and 30.25% (147/486) in Badajoz (Figure 1). However, only 1 out of 25 farms (4%) tested positive for IAV by RT⍰PCR, from which six H1avN2 viruses were obtained in Extremadura. In addition, two H1avN2 viruses and one H3N1 virus were recovered from IAV qPCR positive necropsies samples in this province.

**Figure 1.**
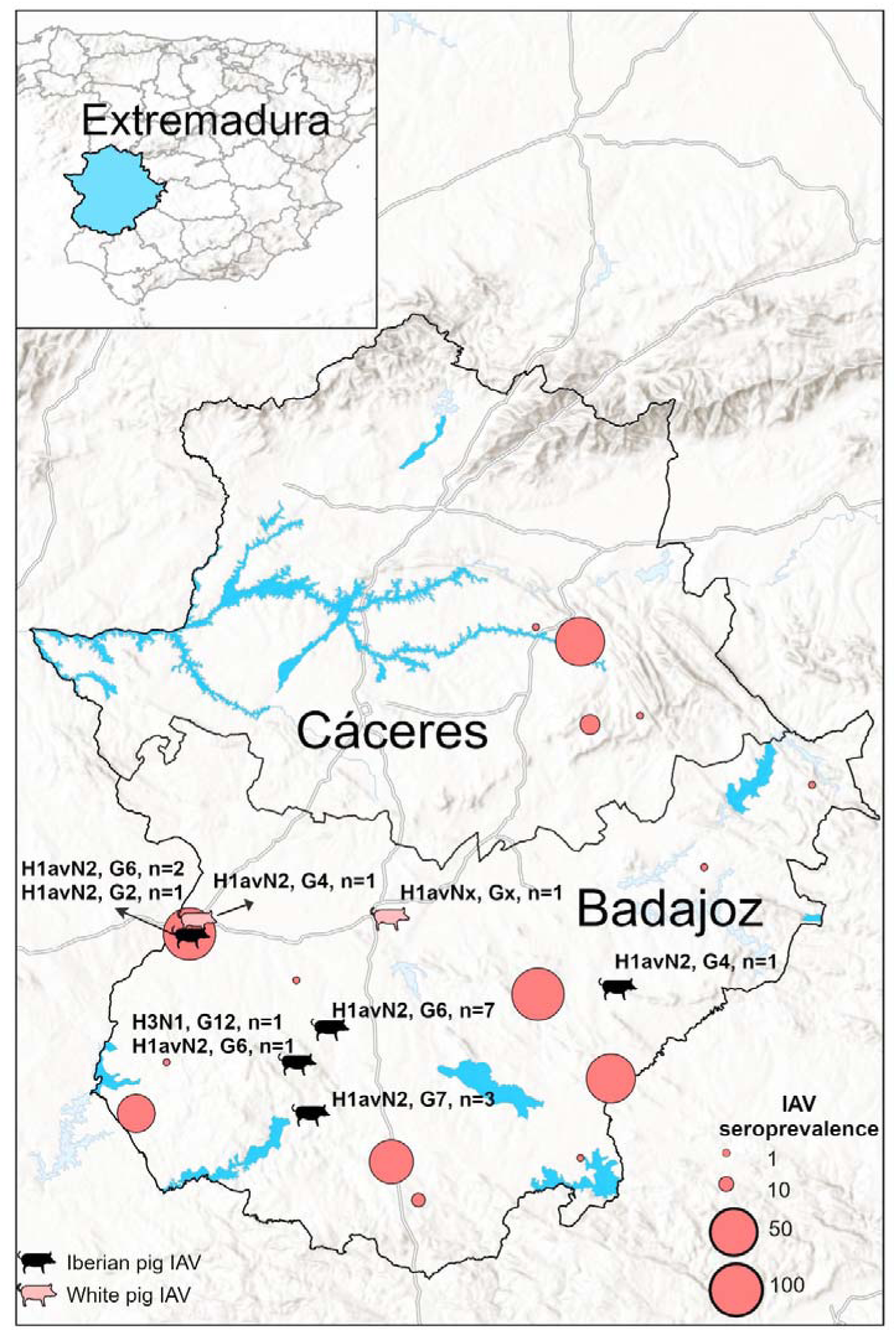
Geographical distribution of the IAV strains detected in commercial Iberian (black) and white pigs (pink) in Extremadura during 2016–2022, and IAV seroprevalence in Iberian pigs in 2020. G: genotypes detected within each subtype, av: avian, x: undetermined, n: number of isolates. Red circles represent the IAV seroprevalence in Iberian pig farms in 2020.

### 3.2 Characterization of SIVs from white pigs (2019–2022)

Of the 108 IAV qPCR-positive samples obtained from Iberian and white pigs during 2019-2022, 57 SIVs were successfully isolated from white pigs. Subtype distribution of those isolates was: 29.8% H1avN2 (17/57), 17.5% H1huN2 (10/57), 12.2% H1av1N1av (7/57), 10.5% H1avN1p (6/57), 8.7% H1pN1p (5/57), 5.2% H1pN1av (3/57), 3.5% H3N1 (2/57), H3N2 (2/57). Five samples could not be fully subtyped.

We identified representatives of all swine H1 lineages currently circulating in pigs in Europe, although most belonged to the Eurasian avian⍰like lineage (EAswH1), which accounted for 55% (32/57) of detections. This was followed by the human seasonal⍰like lineage (HUswH1) at 19% (11/57) and the pandemic⍰derived lineage (PDMswH1) at 17% (10/57). The human seasonal H3 lineage originating in the early 2000s (2000s⍰like H3; clade 2000.3) was detected in 6.9% (4/57) of samples, whereas the human seasonal H3 lineage originating in the 1970s (1970s⍰like H3) lineage was not detected in this study (Figure 2B).

**Figure 2.**
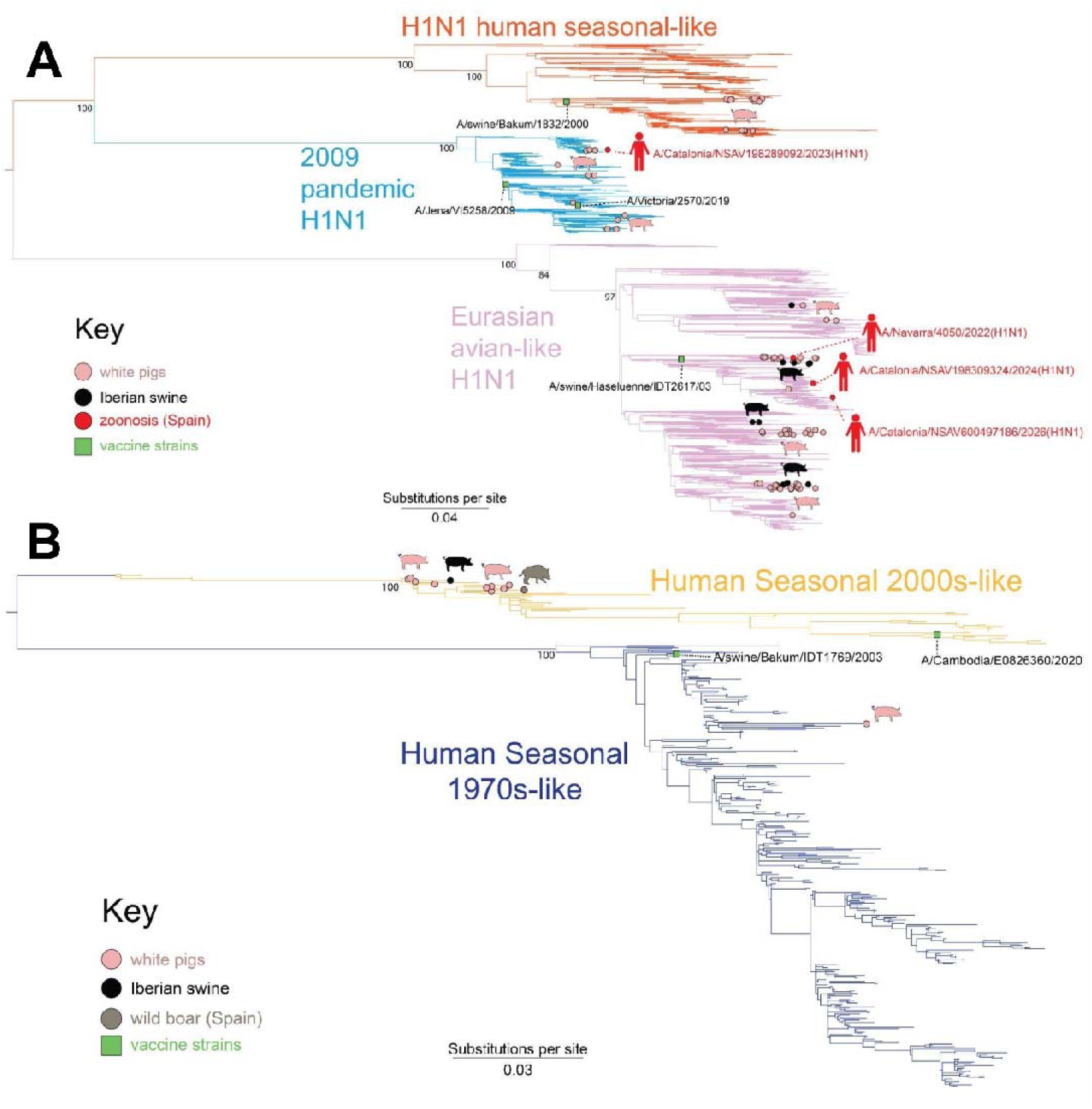
Phylogenetic reconstruction of the H1 (A) and H3 (B) subtypes of European swine and variant IAV. Branches are colored according to IAV lineage. Branch tips representing sequences obtained from commercial white pigs, Iberian pigs, and Spanish zoonotic cases are annotated with pink, black, and red circles, respectively; wild boar sequences (H3 only) are annotated with brown circles. Vaccine strains are shown as green boxes with the corresponding strain name. Spanish variant case strain names are included to annotate zoonotic events. Bootstrap values for major lineages are shown.

The PDMswH1 strains belonged exclusively to clade 1A.3.3.2, while the HUswH1 strains fell within clades 1B.1.2 (6/11) and 1B.1.2.1(5/11). In EAswH1, the most frequently detected subclade was 1C.2.1 (21/32), followed by 1C.2.6 (6/32), 1C.2.4 (3/32) and 1C.2.2 (2/32) at lower frequencies (Figure S1). No 1C.2.3 and 1C.2.5 strains were detected in this study.

The human seasonal N2 lineage from 1970 (1970s-like N2) was the most common, accounting for 50.8% (29/57). The human seasonal N2 lineage from 1990 (1990s-like N2), which had been detected in the previous study in Iberian pig, was not detected in this (Figure 3B). Avian-origin N1 was detected in 17.5% (10/57), and pandemic N1 in 10.5% (6/57) (Figure 3A).

**Figure 3.**
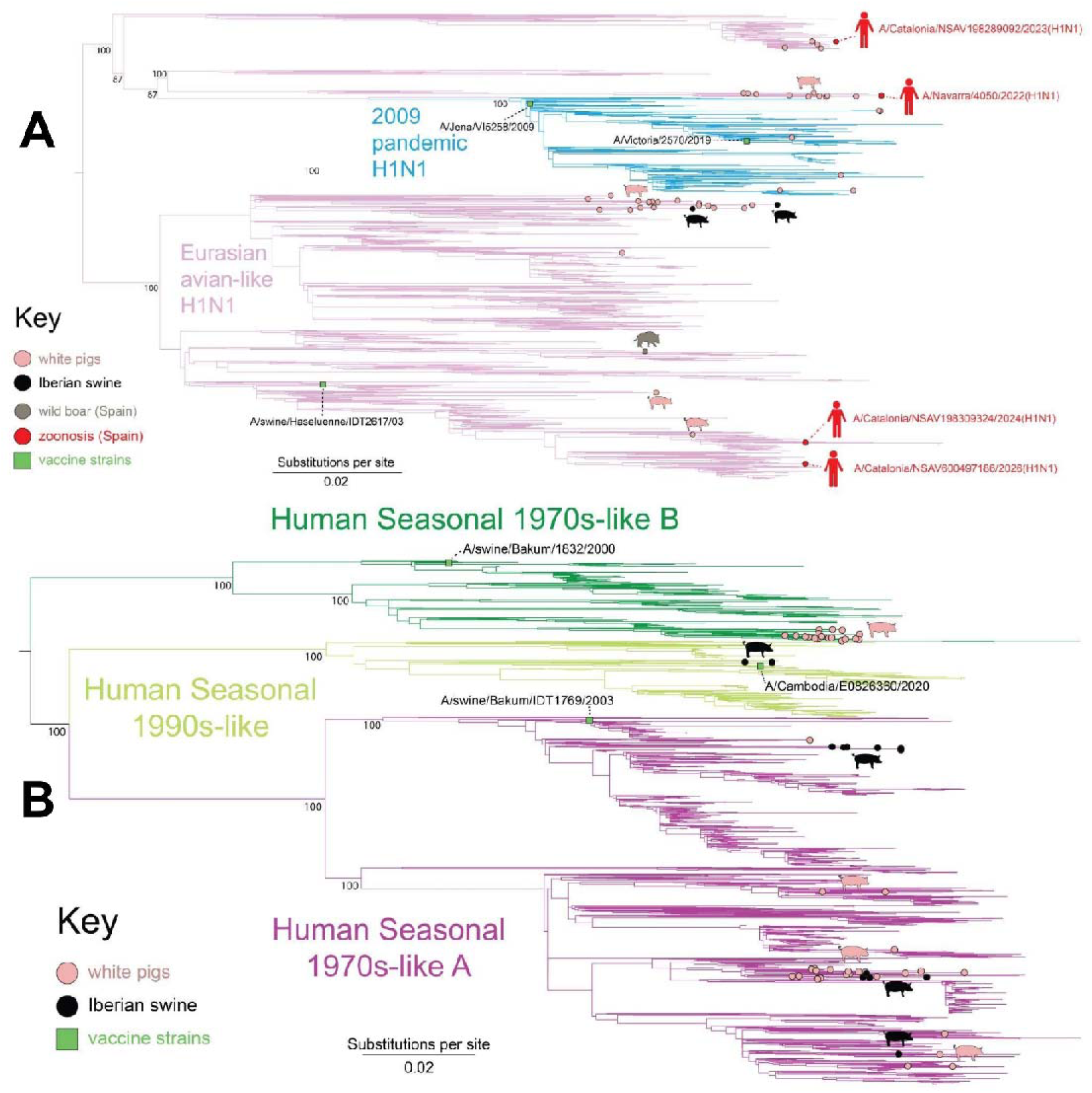
Phylogenetic reconstruction of the N1 (A) and N2 (B) subtypes of European swine and variant IAV. Branches are colored according to IAV lineage. Branch tips representing sequences obtained from commercial white pigs, Iberian pigs, Spanish zoonotic cases, and wild boar (N1 only) are annotated with pink, black, red, and brown circles, respectively. Vaccine strains are shown as green boxes with the corresponding strain name. Spanish variant case strain names are included to annotate zoonotic events. Bootstrap values for major lineages are shown.

### 3.3 Eighteen IAV genotypes identified in swine in Spain (2019–2022)

Of the twelve genotypes identified in our previous study of IAVs in commercial Iberian and white pigs in Spain during 2016-2019 (38), eight were observed again in our most recent investigation (2019–2022) (Figure 4). An additional 11 genotypes (G13-G23) were identified in this last study that were not observed previously. The most frequent genotype in this study was G1.

**Figure 4.**
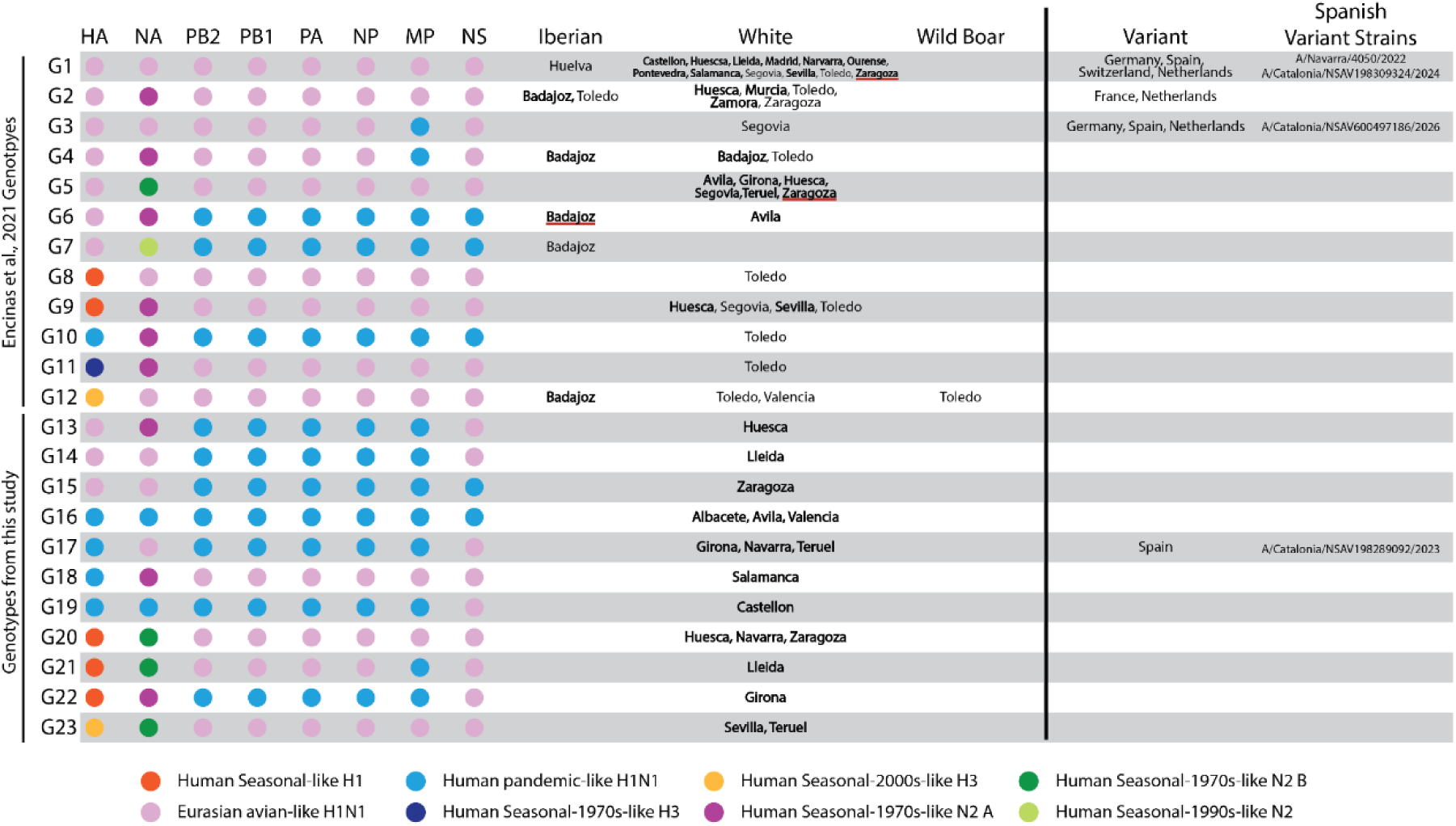
IAV genotypes identified in Iberian and white commercial pigs in Spain from 2016 to 2022. The colors of each circle represent that segment’s lineage for the genotype. The Spanish province in which each genotype was detected within Iberian pigs, commercial white pigs, and wild boar are labeled. Bolded province names signifies that an isolate belonging to the genotype was detected during this current study. Un-bolded province names mean an isolate from that genotype was detected in that province only during our previous study. Provinces underlined in red means isolates from the genotype were found in the province in both studies. Variant cases from Europe and the strain names of variant cases from Spain belonging to the genotypes are included.

G21 and G22 are triple reassortants with surface glycoproteins H1 and N2 of human seasonal origin. G21 has a Eurasian background with a pandemic M segment, whereas G22 has a pandemic background with a Eurasian NS segment (Figure 4, Figure S3). As of April 27, 2026, the H1huN2 G21 and G22 genotypes, as well as H3N1-G12 and H3N2 -G23, had only been detected in pigs in Spain.

The unusual H3N1 “G12” genotype which was first identified in Spanish white pigs in the province of Toledo in 2018, was found in other swine, including a wild boar in that same province in December 2021 (Figure S2) and an Iberian pig in the province of Badajoz in April 2022 (Figure 1). The H3N1-G12 genotype has a genetically distinct H3 that was introduced from humans into European swine in ∼2005, first detected in Germany and Denmark in 2014 and subsequently in Spain in 2018 (A/swine/Spain/143/2018 (H3N2)). In Spain, a genomic reassortment event occurred in which six internal gene segments from the Eurasian avian-like swine lineage were acquired (PB2, PB1, PA, NP, MP, NS), along with a different N2 of human origin that was independently introduced from humans into swine in Europe during the 1970s (N2 lineage “B”), creating a new “G23” genotype in Spain. Through an additional reassortment event, the H3N2-G23 genotype acquired a N1 segment from the Eurasian avian-like swine lineage to create the H3N1-G12 genotype.

### 3.4 Temporal and spatial analysis of the SIV isolates from 2016 to 2022

The SIV diversity found in Iberian pigs (Figure 5A) was limited compared to white pigs (Figure 5B). The main subtype found in Iberian pigs throughout the period was H1avN2. On the other hand, all described SIV subtypes were found in Spanish white pigs, with multiple reassortments that gave rise to new genotypes, which vary in their frequency and time of appearance.

**Figure 5.**
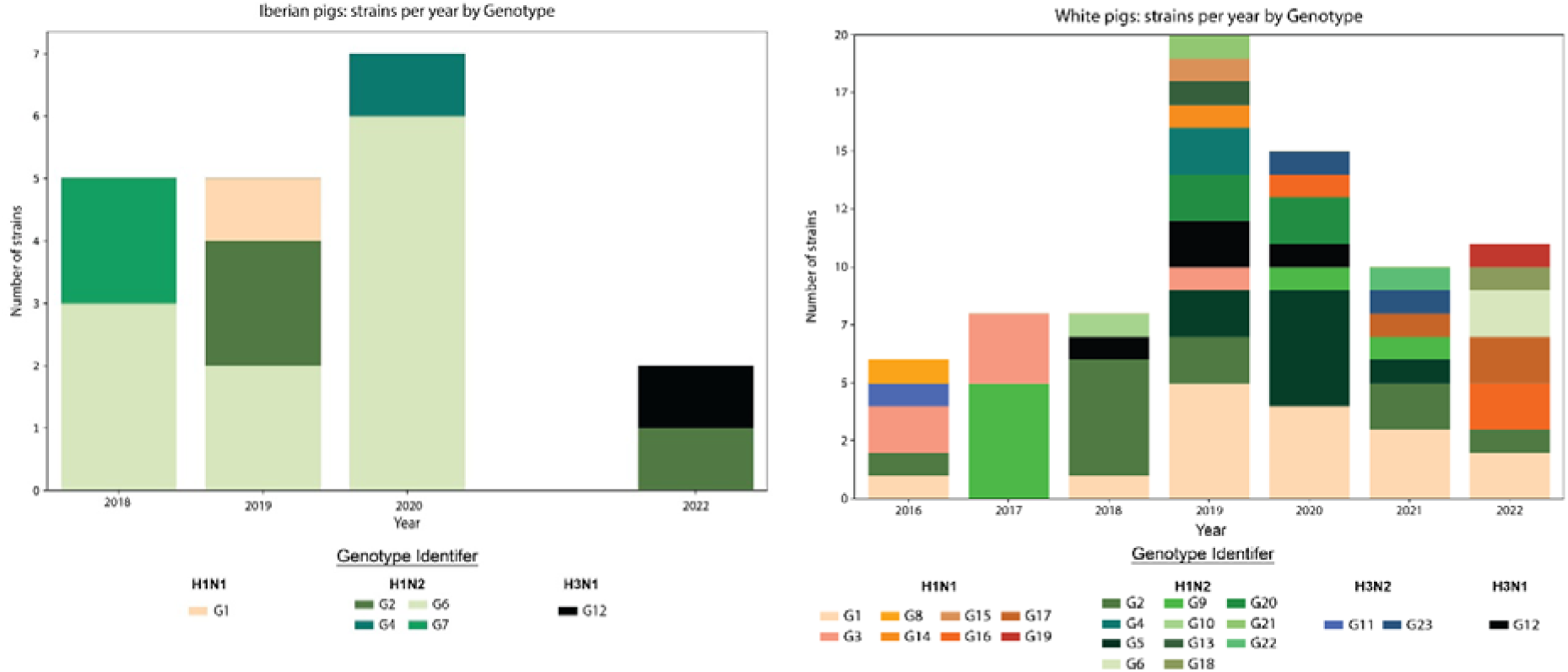
Number of IAV strains detected in Iberian pigs (left) and white pigs (right) each year between 2016-2022 belonging to each genotype.

The G6 genotype, an H1N2A with a pandemic cassette that was only detected in Iberian pigs between 2018 and 2020, appeared in white pigs in 2022 (Figure 5 A, B).

Spatial analysis of SIVs showed that the greatest variety of SIV subtypes and genotypes in white pigs was found in Toledo, followed by Navarra and Zaragoza. However, Huesca showed the highest number of genotypes within a single subtype (Figure 6).

**Figure 6:**
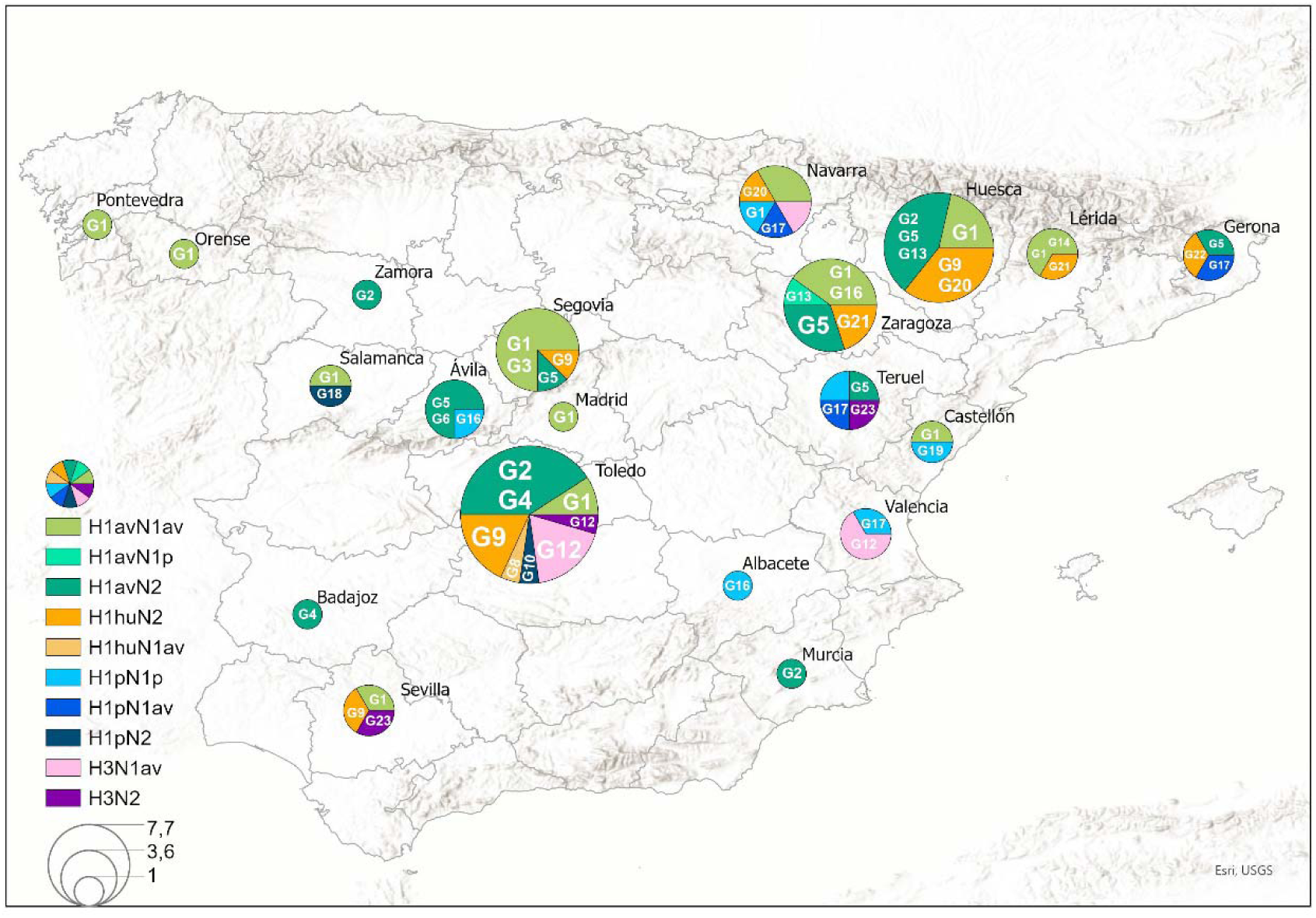
Geographical distribution of the IAV strains detected in commercial white pigs in the 2016-2022 period. G: genotypes detected within each subtype, av: Eurasian lineage, p: pandemic lineage; hu: human lineage. The size of the circles represents the number of samples in each location.

Possible viral transmission between pig breeds was identified in the provinces of Badajoz and Toledo. In Badajoz, H1avN2A(G4) and H1avN2A(G2) were detected in both Iberian and white pigs (Figure 1). Similarly, G2 genotype was also detected in both breeds in Toledo (Figure S2). H3N1 (G12) was detected in white pigs and wild boars in Toledo (Figure S2), and in Iberian pigs in Badajoz (Figure 1). These strains consistenly cluster together with high support values across multiple genomic segments.

### 3.5 Reverse zoonosis of H1pdm09 from humans to swine

Five genotypes detected in Spanish pigs (G10, G16–G19) can be traced back to five independent introductions of the H1N1pdm09 virus from humans into European swine (Figure 7). Eleven viruses identified in Spanish white pigs, as well as one human variant strain (A/Catalonia/NSAV198289092/2023(H1N1)), contained HA segments belonging to the human-origin H1N1pdm09 lineage. In our phylogenetic analysis of the H1N1pdm09 lineage in humans and pigs in Europe from 2009 to 2025, we identified several independent introductions of H1 from humans into pigs across 14 countries. The H1 SIV segments found in Spain trace their ancestry to reverse-zoonotic transmission events originating in Denmark (G17, G18, and G19), Germany (G10), and Spain itself (G16). The G16 genotype evolved twice independently in Spain and is linked to two separate introductions of H1pdm09. A zoonotic IAV case in Spain (A/Catalonia/NSAV198289092/2023) was associated with the G17 genotype. Following its introduction into swine in Italy in 2011, the pdm09 virus reassorted with avian-origin NA and NS segments (Figure 4) and subsequently emerged in Spain and Germany between 2021 and 2022, prior to the zoonotic event reported in Catalonia in 2023.

**Figure 7.**
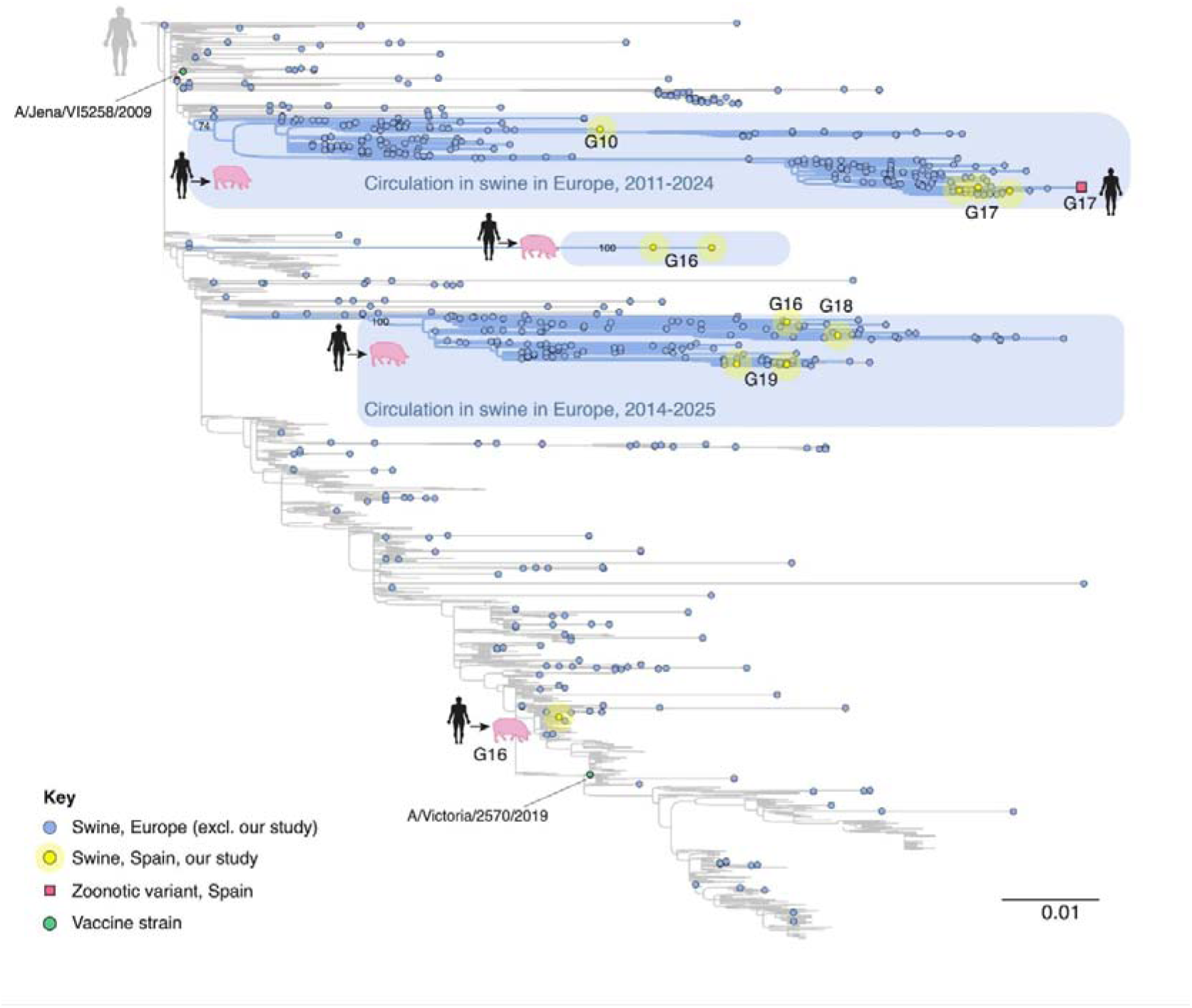
Reverse zoonotic events of H1pdm09 into European swine. Introductions that lead to the emergence of genotypes identified during our study are highlighted in blue. Viruses that belong to genotypes in our study are highlighted in yellow with the corresponding genotype next to it.

### 3.6 Zoonotic infections by the swine-origin H1N1v in Spain, 2022-2026

Four zoonotic cases of swine-origin IAVs belonging to the H1N1 genotype (referred to as “variant” or H1N1v) were detected in Spain during the period 2022-2026 (Figure 8). All four H1N1v zoonotic cases were detected in the northeast region of Spain, where intensive white pig production takes place (48).

**Figure 8.**
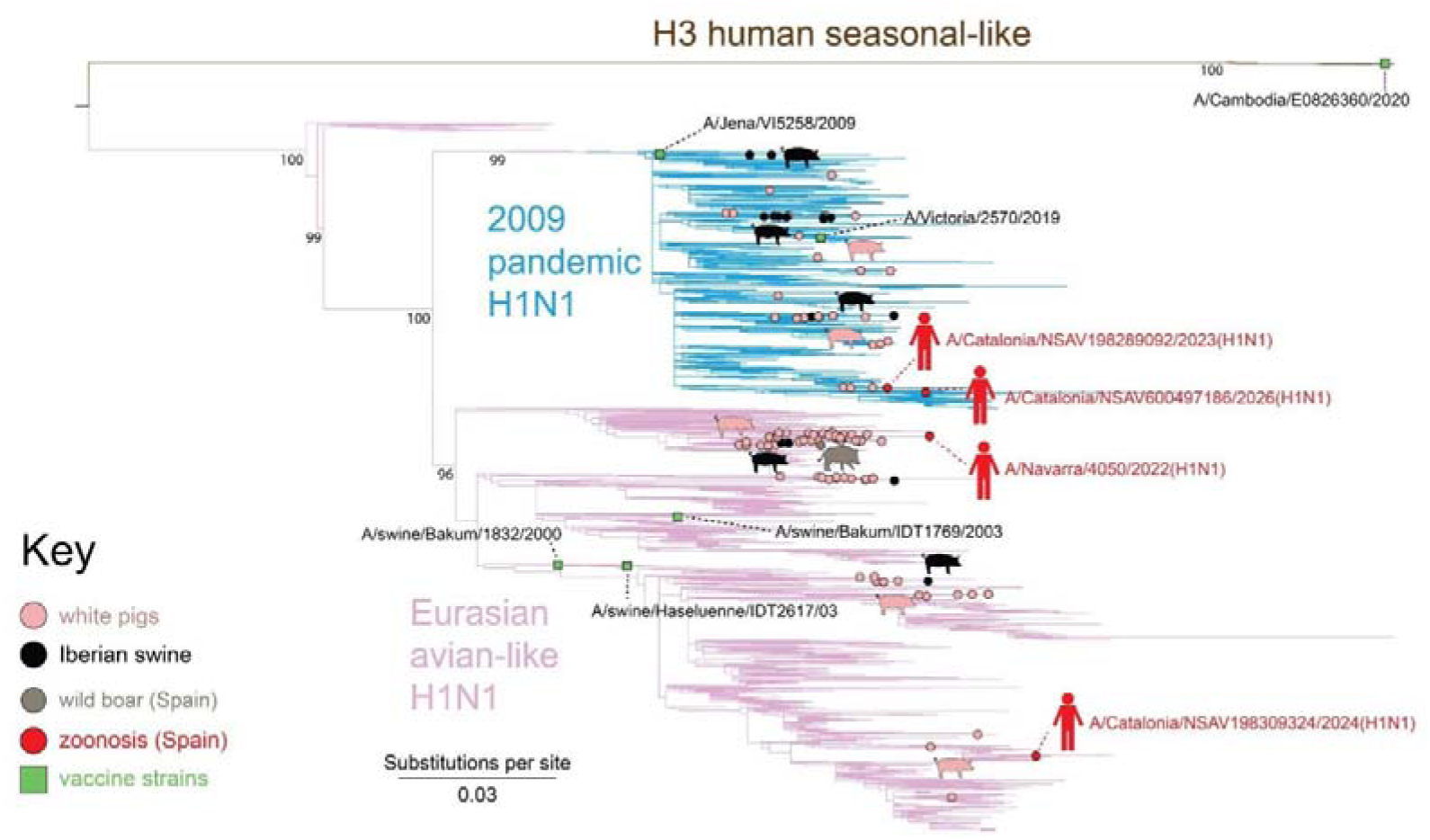
Phylogenetic reconstruction of the MP subtype of European swine and variant IAV. Branches are colored based on IAV lineage. Branch tips representing sequences obtained from commercial white pigs, iberian pigs, wild boar in Spain, and Spanish zoonoses are annotated with pink, black, brown, and red circles, respectively. Vaccine strains are represented as green boxes with the vaccine strain name. Spanish variant case strain names are also included to annotate the zoonotic events. Bootstrap values for lineages are included.

Two zoonotic cases of H1N1v (A/Navarra/4050/2022(H1N1v) and A/Catalonia/NSAV198309324/2024(H1N1v)) were associated with G1 genotype, in which all eight genome segments belong to the Eurasian avian-like H1N1 swine lineage (Figure 4). The G1 genotype was widespread in Spanish white pigs and was identified in more Spanish provinces (n = 12) than any other genotype in our study. This could explain why this genotype, common in white pigs, spilled over into humans on two separate occasions. G1 viruses have circulated in European pigs for decades and have caused zoonotic infections in humans in other countries during the period 2016-2023, including Germany, the Netherlands and Switzerland (Figure S4). A third zoonotic case of H1N1v in Spain (A/Catalonia/NSAV198289092/2023(H1N1v)) was associated with a more recent reassortant genotype G17, which was first detected in pigs in Italy in 2020. G17 viruses have six segments from the human-origin H1N1pdm09 lineage (H1, PB2, PB1, PA, NP, MP) and two segments from the Eurasian avian-like swine lineage (N1 and NS). G17 viruses did not spread as widely in Spanish white pigs as the G1 and G3 genotypes, and were only detected in three provinces (Gerona, Navarra, and Teruel). The G17 swine virus most closely related to A/Catalonia/NSAV198289092/2023(H1N1v) was isolated in this study from a white pig in the province of Gerona in Catalonia region in 2021: A/swine/Spain/17491-1/2022(H1N1). There are no documented cases of the G17 genotype causing zoonotic spillovers in other European countries outside of Spain (Figure 9A). The fourth and most recent zoonotic case of IAV in Spain (A/Catalonia/NSAV600497186/2026(H1N1v)), detected in an 83 years old man with no previous exposure to pigs, was associated with a reassortant G3 genotype in which a pandemic MP segment reassorted into a Eurasian avian-like H1N1 background. G3 viruses were also spilled over to humans in Germany (2022) and the Netherlands (2023) (Figure 9B). In our study, a G3 virus was found in white pigs from Segovia (Figure 6), but it was not closely related to A/Catalonia/NSAV600497186/2026(H1N1v).

**Figure 9.**
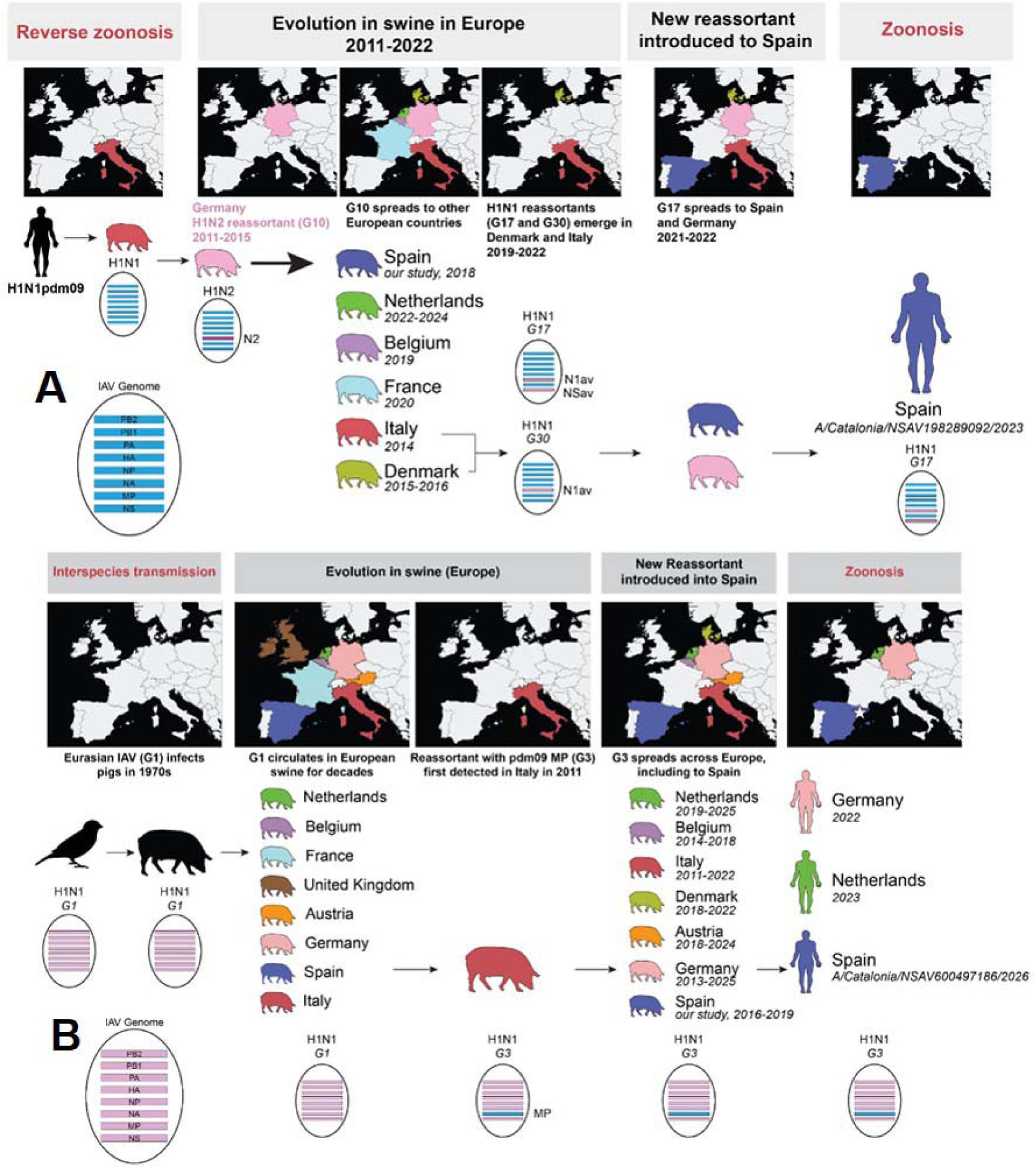
Summary of evolutionary events leading to the appearance of two H1N1v in Spain. Genome segments are ordered by sequence length. **A/Catalonia/NSAV198289092/2023 (H1N1v) (upa.** Following the spread of H1N1pdm09 in humans, a reverse zoonotic event resulted in the transmission of pdm09 into European swine. A H1N1pdm09 strain then reassorted with a human seasonal 1970s-like A NA segment, giving rise to G10. The G10 reassortant was detected in Germany between 2011-2015, before spreading to other European countries in subsequent years. Reassortment of G10 with Eurasian avian NA and NS segments lead to the emergence of the G17 genotype. G17 spread to Germany and Spain in 2021-2022 before the zoonotic transmission in Lerida, Spain in 2023 (A/Catalonia/NSAV198289092/2023 (H1N1)). **B. A/Catalonia/NSAV600497186/2026 (H1N1v).** Genotype 1 (G1) originally transmitted from birds into European swine in the 1970s. G1 circulated in Europe for decades and was detected in multiple countries. In 2011, a reassortant genotype with the pdm09 MP segment (G3) was first detected in Italy. G3 spread to many European countries in the following decade, including Spain. Between 2019 and 2026, three variant cases were associated with G3, one of which occurred in Lerida, Spain (A/Catalonia/NSAV600497186/2026 (H1N1)).

## 4. Discussion

The ecology of IAV in Spain is highly complex, shaped by the coexistence of multiple susceptible host types—white pigs, Iberian pigs, wild boar, and humans—and by diverse production systems (intensive and extensive) operating under closed or open cycles. Many farms are specialized in fattening and routinely import piglets from other countries, mainly the Netherlands, while different production phases may occur on separate farms distributed across distinct regions. In parallel, wild boar move freely across landscapes and interact with multiple environments, and human contact with pigs occurs at numerous points, including farm workers, veterinarians, and transport personnel. Altogether, this combination of host diversity, production structures, animal movements, and human interactions creates a highly dynamic and heterogeneous ecological context for IAV circulation. Within this framework, our study provides seven years (2016–2022) of temporal and spatial data on IAV detection in two pig breeds—white and Iberian —substantially expanding current knowledge of the virus’s ecology in swine in Spain. Moreover, it offers relevant phylogenetic insights into the genetic diversity of circulating IAVs and contributes to clarifying the origins of the human variant cases reported since 2022.

Forty-four percent of the Iberian pig farms tested in 2020 were positive for IAV, with an overall IAV seroprevalence of 28% in Extremadura, slightly lower than that reported for the 2015–2019 period (33.4%). Nine IAVs were successfully isolated from this breed, including two genotypes not previously identified in Iberian pigs (G4 and G12). In contrast, genotypes G6 and G7—both carrying an H1N1pdm09-origin internal cassette and previously detected exclusively in Iberian pigs in the earlier study—showed a different pattern: G7 was not detected in the present sampling, whereas G6 was recently identified in white pigs in 2022 in Ávila.

H3N1 and H3N2 SIV subtypes, carrying a 2000s-like H3 and an Eurasian cassette from human origin, have been consistently detected in domestic pigs and wild boar over the years (Figure 2B, 5A;B). 2000s-like H3 lineage may be also widely disseminated in wild boar, as suggested by a longitudinal surveillance study estimating IAV seroprevalence in this species from 2015 to 2023, which showed that this HA lineage, first detected in 2017 in the Metropolitan Area of Barcelona, has been broadly circulating in these populations since 2021(45). Wild boar may act as a bridging host facilitating the transmission of this lineage to domestic pigs. This may be due to the lack of immunity against this HA lineage in swine populations, given that it was recently introduced from humans into swine. This interpretation is supported by the fact that commercially available SIV vaccines do not include this antigen, and there is no cross-reactivity between 1970s-like and 2000s-like H3, as demonstrated in a serological study testing the cross reactivity of sera from mice immunized with swine influenza vaccines against SIV circulating strains (49).

The 1C.2.1 clade remains by far the most abundant in white pigs; however, some trends can be observed among the less frequent clades. For example, the 1C.2.6 clade, detected in Spain in 2019 (Navarra and Gerona), continued to be identified in subsequent years (2020–2022) across additional provinces, including Huesca, Zaragoza and Pontevedra, suggesting a pattern of ongoing expansion. In contrast, clade 1C.2.2 was detected in only two white pigs in this study, indicating that it may currently be in decline in this breed. Both clades have been detected in three of the H1N1v cases, suggesting a lack of human immunity to these strains. The heterogeneity of human serological responses to EAswH1 IAVs has been demonstrated in several studies. In one study assessing the cross-reactive capacity of sera from individuals immunized with the seasonal influenza vaccine, no antibody response was detected against the strain carrying this HA clade (49) Similar findings were reported in a multi-year longitudinal study, where post-seasonal vaccination hemagglutination inhibition titers against the 1C.2.2 representative virus did not reach the correlate of protection in any year and remained consistently low by microneutralization across all seasons (Lilley et al., 2026).

The high diversity of subtypes and genotypes detected in Toledo, together with the observation that the same genotypes (G2, G12) were found across several pig breeds, suggests that this region may act as a hotspot for interbreed transmission of SIV. Toledo raises both white and Iberian pigs, the latter in smaller numbers, and is located close to Extremadura, the main Iberian pig producing region in Spain, where two genotypes have also been detected in different breeds (G4, G12). In addition, wild boar roam freely across both provinces, providing further opportunities for cross-species and interbreed viral transmission. Further research in this area may help elucidate the exact mechanisms of transmission among different swine populations, thereby enabling the development of targeted containment measures.

Humans are the principal source of pandemic H1N1 introductions into swine populations (50). Since 2009, five independent human⍰to⍰swine introductions of IAV have been documented in Europe, and these viruses have subsequently spread within European pig populations and reassorted with endemic SIV, generating multiple novel genotypes. In our study, there has been a decrease in the number of H1avN1av strains, accompanied by an increase in subtypes carrying an H1/N1pdm component compared with the previous study. Five genomic reassortants have been detected at low frequency. The overall proportion of strains with a complete or nearly complete pandemic cassette in this study was 22%, a value consistent with that reported in a study conducted in Catalonia during 2017–2019.(51). Their genomic constellations include a full pandemic backbone (G16), combined with either avian⍰origin NS (G17, G19) or avian⍰origin N1 (G17), or with a human seasonal 1970s⍰like N2A segment (G10). H1N1pdm09 viruses circulating in swine pose a genuine zoonotic risk, as human immunity to these viruses varies depending on prior vaccination and exposure, and swine⍰adapted H1N1pdm09 lineages likely retain the capacity for efficient human⍰to⍰human transmission (52). Moreover, the independent evolution of this clade in European swine may undermine the effectiveness of human seasonal vaccines, necessitating updates to candidate vaccine viruses, as illustrated by the recent revision prompted by A/Catalonia/NSAV198289092/2023 (H1N1v) (53).

A human variant virus, A/Catalonia/NSAV600497186/2026 (H1N1v), detected in a hospitalized 83-years-old man with no previous exposure to pigs and assigned to G3, carry a pandemic MP segment reassorted into an otherwise fully Eurasian avian-like H1N1 background. This MP segment is phylogenetically related to A/Catalonia/NSAV198289092/2023 (H1N1v) (Figure 8, 9). This case detected recently in Spain was not an isolated event, as two additional G3 H1N1v viruses were identified in Germany (A/Nordrhein-Westfalen/8/2022) and the Netherlands (A/Netherlands/10534/2023), also in individuals without prior pig exposure pigs (54, 55). These latter cases were detected through routine or participatory surveillance systems, and none required hospitalization. In all instances, no further human-to-human transmission has been detected. Studies investigating fitness determinants of IAVs have shown that the phenotype conferred by the pandemic MP segment is critical for viral transmission in humans, and that this M segment increases the transmissibility of swine H3N2 viruses to humans (56). Moreover, the pandemic-origin MP enhances NA activity and transmissibility, suggesting a fitness advantage (56). Altogether, these cases suggest that this genotype, commonly present in European pigs, may be spreading at low, non-detected levels in humans due to improved fitness. In this context, a seroprevalence study among pig⍰farm workers in northeastern Spain would be highly valuable to better assess the prevalence of swine⍰origin IAV in this occupational group, to more accurately determine the true impact of these viruses on the sector, and to better inform biosafety measures.

A key limitation of this study is that we were unable to obtain isolates from all provinces and across all time periods. In addition, the Iberian pig isolates are less numerous and geographically restricted, which limits our ability to draw broader conclusions about their viral diversity and epidemiological dynamics.

## Supporting information

The supplementary material is available in a separate section

## Author Contributions

**Paloma A. Encinas**: conceptualization (lead), investigation (lead), formal analysis (equal), writing – original draft (lead), writing – review and editing (equal). **Brady O’Boyle**: investigation (equal), formal analysis (lead), data curation (lead), writing – original draft (supporting), writing – review and editing (equal), **Alexander Maksaiev**: data curation (equal), formal analysis (equal) **Martha I. Nelson**: investigation (equal), formal analysis (equal), writing – original draft (supporting), writing – review and editing (equal). **Adolfo García-Sastre**: funding acquisition (lead), investigation (equal), project administration (lead), resources (lead), writing – review and editing (equal), **Gustavo del Real**: supervision (lead), investigation (equal), writing – review and editing (equal).

## Acknowledgments

We gratefully acknowledge all data contributors, i.e., the Authors and their Originating laboratories responsible for obtaining the specimens, and their Submitting laboratories for generating the genetic sequence and metadata and sharing via the GISAID Initiative.

## Bibliography

1. Cai D, Shang J, Peng C, Liao H, Shi M, Sun Y. Fast and Accurate Identification of Emerging Viral Reassortment from Genome Sequences. Nucleic Acids Research (2026) 54(6). doi: 10.1093/nar/gkag255.

2. Zhang J, Jiang S, Fang Y, Feng J, Zhang W, Zhang X, et al. Animal Models for Swine Influenza Virus Research: Pathology, Viral Dynamics, and Immune Responses. Viruses (2026) 18(3):344. doi: 10.3390/v18030344.

3. Kong W, Wang F, Dong B, Ou C, Meng D, Liu J, et al. Novel Reassortant Influenza Viruses between Pandemic (H1n1) 2009 and Other Influenza Viruses Pose a Risk to Public Health. Microb Pathog (2015) 89:62–72. Epub 2015/09/08. doi: 10.1016/j.micpath.2015.09.002.

4. Hennig C, Graaf A, Petric PP, Graf L, Schwemmle M, Beer M, et al. Are Pigs Overestimated as a Source of Zoonotic Influenza Viruses? Porcine Health Management (2022) 8(1):30. doi: 10.1186/s40813-022-00274-x.

5. Chauhan RP, Gordon ML. A Systematic Review Analyzing the Prevalence and Circulation of Influenza Viruses in Swine Population Worldwide. Pathogens (2020) 9(5). Epub 20200508. doi: 10.3390/pathogens9050355.

6. Simon G, Larsen LE, Durrwald R, Foni E, Harder T, Van Reeth K, et al. European Surveillance Network for Influenza in Pigs: Surveillance Programs, Diagnostic Tools and Swine Influenza Virus Subtypes Identified in 14 European Countries from 2010 to 2013. PLoS One (2014) 9(12):e115815. Epub 20141226. doi: 10.1371/journal.pone.0115815.

7. Chiapponi C, Prosperi A, Soliani L, De Mattia A, Mescoli A, Torreggiani C, et al. Genetic Diversity and Circulation of Influenza a Viruses in Italian Pig Farms: Insights from Surveillance and Vaccination. Porcine Health Manag (2026) 12(1). Epub 2026/03/14. doi: 10.1186/s40813-026-00498-1.

8. Anderson TK, Chang J, Arendsee ZW, Venkatesh D, Souza CK, Kimble JB, et al. Swine Influenza a Viruses and the Tangled Relationship with Humans. Cold Spring Harb Perspect Med (2021) 11(3). Epub 2020/01/29. doi: 10.1101/cshperspect.a038737.

9. Krog JS, Hjulsager CK, Larsen MA, Larsen LE. Triple-Reassortant Influenza a Virus with H3 of Human Seasonal Origin, Na of Swine Origin, and Internal a(H1n1) Pandemic 2009 Genes Is Established in Danish Pigs. Influenza Other Respir Viruses (2017) 11(3):298–303. Epub 20170321. doi: 10.1111/irv.12451.

10. Krammer F, Barclay WS, Beer M, Brown IH, Cox RJ, de Jong MD, et al. Europe Needs a Sustainably Funded Influenza Research and Response Network. Lancet Infect Dis (2025) 25(4):369–72. Epub 2025/02/21. doi: 10.1016/s1473-3099(25)00068-4.

11. Simon G, Larsen LE, Dürrwald R, Foni E, Harder T, Van Reeth K, et al. European Surveillance Network for Influenza in Pigs: Surveillance Programs, Diagnostic Tools and Swine Influenza Virus Subtypes Identified in 14 European Countries from 2010 to 2013. PLoS One (2014) 9(12):e115815. Epub 2014/12/30. doi: 10.1371/journal.pone.0115815.

12. Ryt-Hansen P, George S, Hjulsager CK, Trebbien R, Krog JS, Ciucani MM, et al. Rapid Surge of Reassortant a(H1n1) Influenza Viruses in Danish Swine and Their Zoonotic Potential. Emerg Microbes Infect (2025) 14(1):2466686. Epub 2025/02/13. doi: 10.1080/22221751.2025.2466686.

13. Richard G, Herve S, Chastagner A, Queguiner S, Beven V, Hirchaud E, et al. Major Change in Swine Influenza Virus Diversity in France Owing to Emergence and Widespread Dissemination of a Newly Introduced H1n2 1c Genotype in 2020. Virus Evol (2025) 11(1):veae112. Epub 2025/01/30. doi: 10.1093/ve/veae112.

14. Klivleyeva N, Saktaganov N, Glebova T, Lukmanova G, Ongarbayeva N, Webby R. Influenza a Viruses in the Swine Population: Ecology and Geographical Distribution. VIRUSES-BASEL (2024) 16(11). doi: 10.3390/v16111728.

15. van der Vries E, Germeraad EA, Kroneman A, Dieste-Pérez L, Eggink D, Willems E, et al. Swine Influenza Virus Surveillance Programme Pilot to Assess the Risk for Animal and Public Health, the Netherlands, 2022 to 2023. Euro Surveill (2025) 30(22). Epub 2025/06/06. doi: 10.2807/1560-7917.Es.2025.30.22.2400664.

16. Lagan P, Hamil M, Cull S, Hanrahan A, Wregor RM, Lemon K. Swine Influenza a Virus Infection Dynamics and Evolution in Intensive Pig Production Systems. Virus Evolution (2024) 10(1). doi: 10.1093/ve/veae017.

17. Lechmann J, Szelecsenyi A, Bruhn S, Harisberger M, Wyler M, Bachofen C, et al. The Swiss National Program for the -Surveillance of Influenza a Viruses in Pigs and Humans: Genetic Variability and Zoonotic Transmissions from 2010 - 2022. Schweiz Arch Tierheilkd (2025) 167(11):600–16. Epub 2025/11/04. doi: 10.17236/sat00466.

18. Rabalski L, Kosinski M, Cybulski P, Stadejek T, Lepek K. Genetic Diversity of Type a Influenza Viruses Found in Swine Herds in Northwestern Poland from 2017 to 2019: The One Health Perspective. Viruses (2023) 15(9). Epub 2023/09/28. doi: 10.3390/v15091893.

19. Kim SW, Gormley A, Jang KB, Duarte ME. - Invited Review - Current Status of Global Pig Production: An Overview and Research Trends. Anim Biosci (2024) 37(4):719–29. Epub 2023/11/10. doi: 10.5713/ab.23.0367.

20. Sanz-Fernández S, Rodríguez-Hernández P, Díaz-Gaona C, Tusell L, Quintanilla R, Rodríguez-Estévez V. Evolution of Sow Productivity and Evaluation Parameters: Spanish Farms as a Benchmark. Vet Sci (2024) 11(12). Epub 2024/12/27. doi: 10.3390/vetsci11120626.

21. Sylven KR, Jacobson M, Schwarz L, Zohari S. Reverse Zoonotic Transmission of Human Seasonal Influenza to a Pig Herd in Sweden. Tierarztl Prax Ausg G Grosstiere Nutztiere (2024) 52(5):296–303. Epub 2024/10/25. doi: 10.1055/a-2410-1530.

22. Jore S, Tønnessen R, Grøntvedt CA, Rydland K, Hauge AG, Hungnes O, et al. A Retrospective Cross-Sectional Study (2009–2023): Exploring Influenza a(H1n1)Pdm09 Antibody Time Series in Humans and Swine and Vaccine Coverage in Two Target Groups. Zoonoses and Public Health (2026) 73(4):336–47. doi: 10.1111/zph.70051.

23. Muthappan S, Abdulkader RS, Mohd G, Beryl Lydia J, Priya J, Salvankar A, et al. Risk of Swine Influenza Virus Spillover at the Human-Swine Interface - a Scoping Review. Int J Public Health (2025) 70:1608380. Epub 2025/10/06. doi: 10.3389/ijph.2025.1608380.

24. Forgie SE, Keenliside J, Wilkinson C, Webby R, Lu P, Sorensen O, et al. Swine Outbreak of Pandemic Influenza a Virus on a Canadian Research Farm Supports Human-to-Swine Transmission. Clin Infect Dis (2011) 52(1):10–8. Epub 2010/12/15. doi: 10.1093/cid/ciq030.

25. Ministry of Agriculture, Fisheries and Food. Resultados De Las Encuestas De Ganado Porcino Noviembre 2024. Ministry of Agriculture, Fisheries and Food (2024).

26. Ruiz-Rodriguez C, Blanco-Aguiar JA, Fernandez-Lopez J, Acevedo P, Montoro V, Illanas S, et al. A Methodological Framework to Characterize the Wildlife-Livestock Interface: The Case of Wild Boar in Mainland Spain. Prev Vet Med (2024) 230:106280. Epub 2024/07/26. doi: 10.1016/j.prevetmed.2024.106280.

27. Triguero-Ocaña R, Laguna E, Jiménez-Ruiz S, Fernández-López J, García-Bocanegra I, Barasona J, et al. The Wildlife-Livestock Interface on Extensive Free-Ranging Pig Farms in Central Spain During the “Montanera” Period. Transbound Emerg Dis (2021) 68(4):2066–78. Epub 2020/09/27. doi: 10.1111/tbed.13854.

28. Zeller MA, Carnevale de Almeida Moraes D, Ciacci Zanella G, Souza CK, Anderson TK, Baker AL, et al. Reverse Zoonosis of the 2022–2023 Human Seasonal H3n2 Detected in Swine. npj Viruses (2024) 2(1):27. doi: 10.1038/s44298-024-00042-4.

29. Soliani L, Mescoli A, Zanni I, Baioni L, Alborali G, Moreno A, et al. Human-Derived H3n2 Influenza a Viruses Detected in Pigs in Northern Italy. Viruses (2025) 17(9). Epub 2025/09/27. doi: 10.3390/v17091171.

30. Nelson MI, Wentworth DE, Culhane MR, Vincent AL, Viboud C, LaPointe MP, et al. Introductions and Evolution of Human-Origin Seasonal Influenza a Viruses in Multinational Swine Populations. Journal of Virology (2014) 88(17):10110–9. doi: doi:10.1128/jvi.01080-14.

31. Klivleyeva N, Glebova T, Saktaganov N, Webby R. Cases of Interspecies Transmission of Influenza a Virus from Swine to Humans. Vet Sci (2025) 12(9). Epub 2025/09/27. doi: 10.3390/vetsci12090873.

32. Eggink D, Kroneman A, Dingemans J, Goderski G, van den Brink S, Bagheri M, et al. Human Infections with Eurasian Avian-Like Swine Influenza Virus Detected by Coincidence Via Routine Respiratory Surveillance Systems, the Netherlands, 2020 to 2023. Euro Surveill (2025) 30(19). Epub 2025/05/16. doi: 10.2807/1560-7917.Es.2025.30.19.2400662.

33. Heider A, Wedde M, Weinheimer V, Döllinger S, Monazahian M, Dürrwald R, et al. Characteristics of Two Zoonotic Swine Influenza a(H1n1) Viruses Isolated in Germany from Diseased Patients. Int J Med Microbiol (2024) 314:151609. Epub 2024/01/29. doi: 10.1016/j.ijmm.2024.151609.

34. Cogdale J, Kele B, Myers R, Harvey R, Lofts A, Mikaiel T, et al. A Case of Swine Influenza a(H1n2)V in England, November 2023. Euro Surveill (2024) 29(3). Epub 2024/01/19. doi: 10.2807/1560-7917.Es.2024.29.3.2400002.

35. Andersen KM, Vestergaard LS, Nissen JN, George SJ, Ryt-Hansen P, Hjulsager CK, et al. Severe Human Case of Zoonotic Infection with Swine-Origin Influenza a Virus, Denmark, 2021. Emerg Infect Dis (2022) 28(12):2561-4. Epub 2022/11/24. doi: 10.3201/eid2812.220935.

36. World Health Organization. Reports on Epidemic Outbreaks; Variant of the Influenza a(H1n1) Virus – Spain. (2024).

37. Markin A, Ciacci Zanella G, Arendsee ZW, Zhang J, Krueger KM, Gauger PC, et al. Reverse-Zoonoses of 2009 H1n1 Pandemic Influenza a Viruses and Evolution in United States Swine Results in Viruses with Zoonotic Potential. PLoS Pathog (2023) 19(7):e1011476. Epub 20230727. doi: 10.1371/journal.ppat.1011476.

38. Encinas P, Del Real G, Dutta J, Khan Z, van Bakel H, Del Burgo MAM, et al. Evolution of Influenza a Virus in Intensive and Free-Range Swine Farms in Spain. Virus Evol (2022) 7(2):veab099. Epub 20211130. doi: 10.1093/ve/veab099.

39. World Health Organization (WHO). Manual for the Laboratory Diagnosis and Virological Surveillance of Influenza. (2011).

40. Zhou B, Donnelly ME, Scholes DT, St George K, Hatta M, Kawaoka Y, et al. Single-Reaction Genomic Amplification Accelerates Sequencing and Vaccine Production for Classical and Swine Origin Human Influenza a Viruses. J Virol (2009) 83(19):10309–13. Epub 2009/07/17. doi: 10.1128/jvi.01109-09.

41. Haas BJ, Papanicolaou A, Yassour M, Grabherr M, Blood PD, Bowden J, et al. De Novo Transcript Sequence Reconstruction from Rna-Seq Using the Trinity Platform for Reference Generation and Analysis. Nat Protoc (2013) 8(8):1494–512. Epub 2013/07/13. doi: 10.1038/nprot.2013.084.

42. Minh BQ, Schmidt HA, Chernomor O, Schrempf D, Woodhams MD, von Haeseler A, et al. Iq-Tree 2: New Models and Efficient Methods for Phylogenetic Inference in the Genomic Era. Molecular Biology and Evolution (2020) 37(5):1530–4. doi: 10.1093/molbev/msaa015.

43. Rambaut A, Lam TT, Max Carvalho L, Pybus OG. Exploring the Temporal Structure of Heterochronous Sequences Using Tempest (Formerly Path-O-Gen). Virus Evol (2016) 2(1):vew007. Epub 2016/10/25. doi: 10.1093/ve/vew007.

44. Rambaut A. Figtree. University of Edinburgh: Institute of Evolutionary Biology (2010).

45. Encinas P, Nogales A, Escribano-Romero E, Del Burgo MAM, Lopez-Olvera JR, Granados JE, et al. Longitudinal Surveillance of Influenza a Virus Exposure in Wild Boars (Sus Scrofa) in Spain (2015-2023): Serologic and Virologic Evidence of Subtype Infections and H5n1 Spillover Risk. Zoonoses Public Health (2026) 73(3):191–204. Epub 2026/02/11. doi: 10.1111/zph.70040.

46. Olson RD, Assaf R, Brettin T, Conrad N, Cucinell C, Davis JJ, et al. Introducing the Bacterial and Viral Bioinformatics Resource Center (Bv-Brc): A Resource Combining Patric, Ird and Vipr. Nucleic Acids Res (2023) 51(D1):D678–d89. Epub 2022/11/10. doi: 10.1093/nar/gkac1003.

47. Katoh K, Standley DM. Mafft Multiple Sequence Alignment Software Version 7: Improvements in Performance and Usability. Mol Biol Evol (2013) 30(4):772–80. Epub 2013/01/19. doi: 10.1093/molbev/mst010.

48. Eurostat. Agriculture Statistics at Regional Level. (2025).

49. Encinas P, Lalueza-Blanco A, Garcia-Vaquero AI, Nelson MI, Garcia-Sastre A, Del Real G. Seasonal Influenza Vaccine 2021/2022 Provides Limited Cross Reactivity against Contemporary Swine Influenza a Virus Strains in Spain. Influenza Other Respir Viruses (2025) 19(12):e70158. Epub 2025/12/23. doi: 10.1111/irv.70158.

50. Ciacci Zanella G, Markin A, Neveau Thomas M, Snyder Celeste A, Souza Carine K, Arruda B, et al. Transmission and Pathologic Findings of Divergent Human Seasonal H1n1pdm09 Influenza a Viruses Following Spillover into Pigs in the United States. Influenza and Other Respiratory Viruses (2025) 19(7):e70128. doi: 10.1111/irv.70128.

51. Sosa Portugal S, Cortey M, Tello M, Casanovas C, Mesonero-Escuredo S, Barrabes S, et al. Diversity of Influenza a Viruses Retrieved from Respiratory Disease Outbreaks and Subclinically Infected Herds in Spain (2017-2019). Transbound Emerg Dis (2021) 68(2):519–30. Epub 2020/07/04. doi: 10.1111/tbed.13709.

52. Cook PW, Stark T, Jones J, Kondor R, Zanders N, Benfer J, et al. Detection and Characterization of Swine Origin Influenza a(H1n1) Pandemic 2009 Viruses in Humans Following Zoonotic Transmission. J Virol (2020) 95(2). Epub 20201222. doi: 10.1128/JVI.01066-20.

53. World Health Organization (WHO). Summary of Status of Development and Availability of Variant1 Influenza a(H1) Candidate Vaccine Viruses and Potency Testing Reagents. (2026).

54. World Health Organization (WHO). Influenza a H1n1 - Germany. (2022).

55. World Health Organization (WHO). Influenza a (H1n1) Variant Virus - the Netherlands. (2023).

56. Griffin EF, Tompkins SM. Fitness Determinants of Influenza a Viruses. Viruses (2023) 15(9). Epub 2023/09/28. doi: 10.3390/v15091959.

