## Supplementary material for "Zoonotic infections and genomic evolution of novel reassortant swine influenza A viruses in Spain"

**Table S1 GenBank Accession Numbers**

| **STRAIN NAME** | **PB2** | **PB1** | **PA** | **HA** | **NP** | **NA** | **MP** | **NS** |
| --- | --- | --- | --- | --- | --- | --- | --- | --- |
| A/swine/Spain/50001-3/2019 | MW848751.1 | MW848747.1 |  | MW848749.1 | MW848750.1 | MW848752.1 | MW848748.1 | MW848746.1 |
| A/swine/Spain/50001-2/2019 | MW848645.1 | MW848639.1 | MW848638.1 | MW848643.1 | MW848642.1 | MW848641.1 | MW848640.1 | MW848644.1 |
| A/swine/Spain/000-1/2019 | MW848730.1 | MW848733.1 | MW848731.1 | MW848735.1 | MW848732.1 | MW848734.1 | MW848736.1 | MW848737.1 |
| A/swine/Spain/31001-2/2019 | MW848686.1 |  | MW848685.1 | MW848689.1 | MW848684.1 | MW848683.1 | MW848688.1 | MW848687.1 |
| A/swine/Spain/06001-3/2019 | MW848779.1 | MW848776.1 | MW848775.1 | MW848781.1 | MW848777.1 | MW848778.1 | MW848780.1 | MW848774.1 |
| A/swine/Spain/50001-4/2019 | MW848681.1 | MW851890.1 | MW848678.1 |  | MW848679.1 | MW848677. | MW848682.1 | MW848680.1 |
| A/swine/Spain/50001-6/2019 | MW848383.1 | MW848386.1 | MW848381.1 | MW848380.1 | MW848385.1 | MW848387.1 | MW848384.1 | MW848382.1 |
| A/swine/Spain/25001-1/2019 | MW848723.1 | MW851851.1 | MW848726.1 | MW848728.1 | MW848724.1 | MW848725.1 | MW848727.1 | MW848729.1 |
| A/swine/Spain/25001-2/2019 | MW848669.1 | MW848665.1 | MW848667.1 | MW848663.1 | MW848662.1 | MW848668.1 | MW848666.1 | MW848664.1 |
| A/swine/Spain/50001-5/2019 | MW848661.1 | MW848657.1 | MW848660.1 | MW848655.1 | MW848658.1 | MW848656.1 | MW848654.1 | MW848659.1 |
| A/swine/Spain/17001-1/2019 |  | MW848708.1 | MW848705.1 | MW848707.1 | MW848706.1 |  | MW848704.1 | MW848703.1 |
| A/swine/Spain/25001-3/2019 | MW848652.1 | MW848650.1 | MW848651.1 | MW848648.1 | MW848647.1 | MW848649.1 | MW848646.1 | MW848653.1 |
| A/swine/Spain/25001-4/2019 | MW848713.1 | MW851852.1 | MW848709.1 | MW848714.1 | MW848710.1 | MW848711.1 | MW848712.1 | MW848715.1 |
| A/swine/Spain/6370-7/2020 | MW848767.1 | MW851848.1 | MW848772.1 | MW848773.1 | MW848771.1 | MW848768.1 | MW848770.1 | MW848769.1 |
| A/swine/Spain/6370-6/2020 | MW848696.1 | MW851853.1 | MW848697.1 | MW848699.1 | MW848698.1 | MW848700.1 | MW848702.1 | MW848701.1 |
| A/swine/Spain/6370-8/2020 | PQ107519 | PQ107520 | PQ107521 | PQ107522 | PQ107523 | PQ107524 | PQ107525 | PQ107526 |
| A/swine/Spain/6370-9/2020 | PQ107529 | PQ107530 | PQ107531 | PQ107532 | PQ107533 | PQ107534 | PQ107535 | PQ107536 |
| A/swine/Spain/6370-10/2020 | PQ107537 | PQ107538 | PQ107539 | PQ107540 | PQ107541 | PQ107542 | PQ107543 | PQ107544 |
| A/swine/Spain/6370-11/2020 | PQ107554 | PQ107555 | PQ107556 | PQ107557 | PQ107558 | PQ107559 | PQ107560 | PQ107561 |
| A/swine/Spain/46314-1/2020 | MW848675.1 |  | MW848674.1 | MW848671.1 | MW848676.1 | MW848672.1 | MW848670.1 | MW848673.1 |
| A/swine/Spain/22000-1/2020 | MW848743.1 | MW848744.1 | MW848740.1 | MW848738.1 | MW848742.1 | MW848745.1 | MW848739.1 | MW848741.1 |
| A/swine/Spain/44490-1/2020 | MW848628.1 | MW848622.1 | MW848626.1 | MW848623.1 | MW848624.1 | MW848629.1 | MW848625.1 | MW848627.1 |
| A/swine/Spain/22251-1/2020 |  | MW848719.1 | MW848716.1 | MW848721.1 | MW848718.1 | MW848717.1 | MW848720.1 | MW848722.1 |
| A/swine/Spain/46314-2/2020 | MW848634.1 | MW848633.1 | MW848630.1 | MW848635.1 | MW848631.1 | MW848637.1 | MW848632.1 | MW848636.1 |
| A/swine/Spain/50800-1/2020 | MW848755.1 |  | MW848753.1 | MW848758.1 | MW848757.1 | MW848759.1 | MW848756.1 | MW848754.1 |
| A/swine/Spain/44579-1/2020 | MW848766.1 | MW848763.1 | ON716287.1 | MW848761.1 | MW848760.1 | MW848765.1 | MW848762.1 | MW848764.1 |
| A/swine/Spain/22269-1/2020 | PP338874 | PP338875 |  | PP338876 | PP338877 | PP338878 | PP338879 | PP338880 |
| A/swine/Spain/31310-1/2020 | PP330401 | PP330402 | PP330403 | PP330404 | PP330405 | PP330406 | PP330407 | PP330408 |
| A/swine/Spain/22070-1/2020 | PP330414 | PP330415 |  | PP330416 | PP330417 | PP330418 | PP330419 | PP330420 |
| A/swine/Spain/50070-1/2020 | PP330721 | PP330722 | PP330723 | PP330724 | PP330725 | PP330726 | PP330727 | PP330728 |
| A/swine/Spain/32070-1/2020 | PP330730 | PP330731 | PP330732 | PP330733 | PP330734 | PP330735 | PP330736 | PP330737 |
| A/swine/Spain/22270-1/2020 | PP330763 | PP330764 | PP330765 | PP330766 | PP330767 | PP330768 | PP330769 | PP330770 |
| A/swine/Spain/40217-1/2020 | PP331797 | PP331798 | PP331799 | PP331800 | PP331801 | PP331802 | PP331803 | PP331804 |
| A/swine/Spain/05165-1/2020 | PP331788 | PP331789 | PP331790 | PP331791 | PP331792 | PP331793 | PP331794 | PP331795 |
| A/swine/Spain/22270-2/2020 | PP335236 | PP335237 | PP335238 | PP335239 | PP335240 | PP335241 | PP335242 | PP335243 |
| A/swine/Spain/25211-1/2020 | PP335491 | PP335492 | PP335493 | PP335494 | PP335495 | PP335496 | PP335497 | PP335498 |
| A/swine/Spain/05530-1/2020 | PP335504 | PP335505 | PP335506 | PP335507 | PP335508 | PP335509 | PP335510 | PP335511 |
| A/swine/Spain/06800-1/2020 | PQ580717 | PQ580718 | PQ580719 | PQ580720 | PQ580721 |  | PQ580722 | PQ580723 |
| A/swine/Spain/22220-1/2020 | PQ585394 | PQ585395 | PQ585396 | PQ585397 | PQ585398 | PQ585399 | PQ585400 | PQ585401 |
| A/swine/Spain/28000-1/2021 | PP338223 | PP338224 | PP338225 | PP338226 | PP338227 | PP338228 | PP338229 | PP338230 |
| A/swine/Spain/06070-1/2020 | PP338238 | PP338239 | PP338240 | PP338241 | PP338242 | PP338243 | PP338244 | PP338245 |
| A/swine/Spain/17473-1/2021 | PP338247 | PP338248 | PP338249 | PP338250 | PP338251 | PP338252 | PP338253 | PP338254 |
| A/swine/Spain/22253-1/2020 | PP338272 | PP338273 | PP338274 | PP338275 | PP338276 | PP338277 | PP338278 | PP338279 |
| A/swine/Spain/41566-1/2021 | PP338281 | PP338282 | PP338283 | PP338284 | PP338285 | PP338286 | PP338287 | PP338288 |
| A/swine/Spain/41410-1/2021 | PP338503 | PP338504 | PP338505 | PP338506 | PP338507 | PP338508 | PP338509 | PP338510 |
| A/swine/Spain/22192-1/2021 | PP338511 | PP338512 | PP338513 | PP338514 | PP338515 | PP338516 | PP338517 | PP338518 |
| A/swine/Spain/44593-1/2022 |  | PP338519 | PP338520 | PP338521 | PP338522 | PP338523 | PP338524 | PP338525 |
| A/swine/Spain/30870-1/2021 | PP338530 | PP338531 | PP338532 | PP338533 | PP338534 | PP338535 | PP338536 | PP338537 |
| A/swine/Spain/12312-1/2021 | PP338682 | PP338683 | PP338684 | PP338685 | PP338686 | PP338687 | PP338688 | PP338689 |
| A/swine/Spain/22260-1/2021 | PP338713 | PP338714 | PP338715 | PP338716 | PP338717 | PP338718 | PP338719 | PP338720 |
| A/swine/Spain/17483-1/2021 | PQ107546 | PQ107547 | PQ107548 | PQ107549 | PQ107550 | PQ107551 | PQ107552 | PQ107553 |
| A/swine/Spain/44394-1/2021 | PQ107577 | PQ107578 | PQ107579 | PQ107580 | PQ107581 | PQ107582 | PQ107583 | PQ107584 |
| A/swine/Spain/05165-2/2022 | PQ107585 | PQ107586 | PQ107587 | PQ107588 | PQ107589 | PQ107590 | PQ107591 | PQ107592 |
| A/swine/Spain/37500-1/2022 | PQ107595 | PQ107596 | PQ107597 | PQ107598 | PQ107599 | PQ107600 | PQ107601 | PQ107602 |
| A/swine/Spain/6176-2/2022 | PQ107608 | PQ107609 | PQ107610 | PQ107611 | PQ107612 | PQ107613 | PQ107614 | PQ107615 |
| A/swine/Spain/12510-1/2022 | PQ107616 | PQ107617 | PQ107618 | PQ107619 | PQ107620 | PQ107621 | PQ107622 | PQ107623 |
| A/swine/Spain/49610-1/2022 | PQ107624 | PQ107625 | PQ107626 | PQ107627 | PQ107628 | PQ107629 | PQ107630 | PQ107631 |
| A/swine/Spain/2247-1/2022 | PQ107637 | PQ107638 | PQ107639 | PQ107640 | PQ107641 | PQ107642 | PQ107643 | PQ107644 |
| A/swine/Spain/31510-1/2022 | PQ107645 | PQ107646 | PQ107647 | PQ107648 | PQ107649 | PQ107650 | PQ107651 | PQ107652 |
| A/swine/Spain/06001-4/2022 | PQ108487 | PQ108488 | PQ108489 | PQ108490 | PQ108491 | PQ108492 | PQ108493 | PQ108494 |
| A/swine/Spain/17491-1/2022 | PQ111069 | PQ111070 | PQ111071 | PQ111072 | PQ111073 | PQ111074 | PQ111075 | PQ111075 |
| A/swine/Spain/22210-1/2022 | PQ107666 | PQ107667 | PQ107668 |  | PQ107669 | PQ107670 | PQ107671 | PQ107672 |
| A/swine/Spain/46892-1/2022 | PQ107658 | PQ107659 | PQ107660 | PQ107661 | PQ107662 | PQ107663 | PQ107664 | PQ107665 |
| A/swine/Spain/5296-1/2022 | PQ585327 | PQ585328 | PQ585329 | PQ585330 | PQ585331 | PQ585332 | PQ585333 | PQ585334 |
| A/swine/Spain/45790-1/2022 | PQ107769 |  | PQ107770 | PQ107771 | PQ107772 | PQ107773 | PQ107774 | PQ107775 |
| A/swine/Spain/36520-1/2022 | PQ585340 | PQ585341 | PQ585342 | PQ585343 | PQ585344 | PQ585345 | PQ585346 | PQ585347 |
| A/swine/Spain/50700-1/2022 | PQ107776 | PQ107777 | PQ107778 | PQ107779 | PQ107780 | PQ107781 | PQ107782 | PQ107783 |

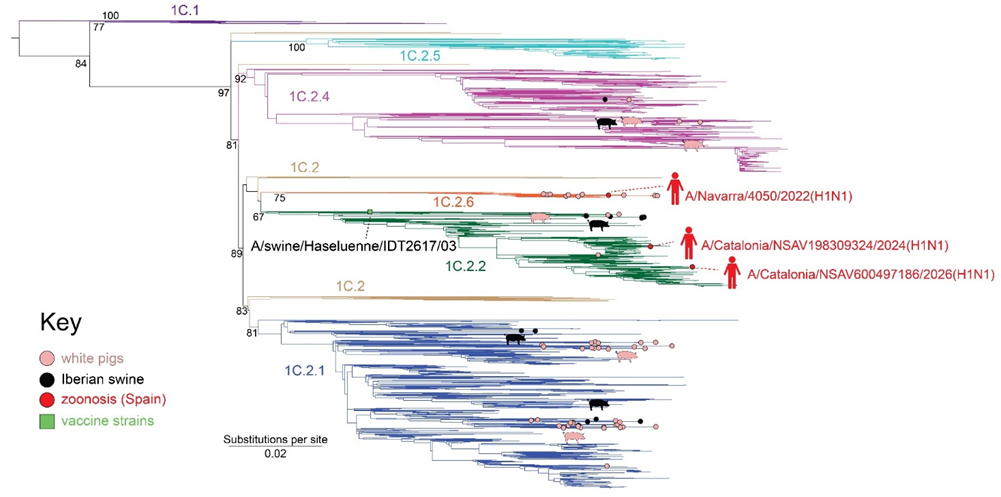

**Figure S1 Phylogenetic reconstruction of human and swine European H1 Eurasian Avian-like sequences from 2000-2022**. Branches are colored based on IAV Eurasian Avian-like lineage. Branch tips representing sequences obtained from commercial white pigs, Iberian pigs, and Spanish zoonoses are annotated with pink, black, and red circles, respectively. Vaccine strains are represented as green boxes with the vaccine strain name. Spanish variant case strain names are also included to annotate the zoonotic events. Bootstrap values for lineages are included.

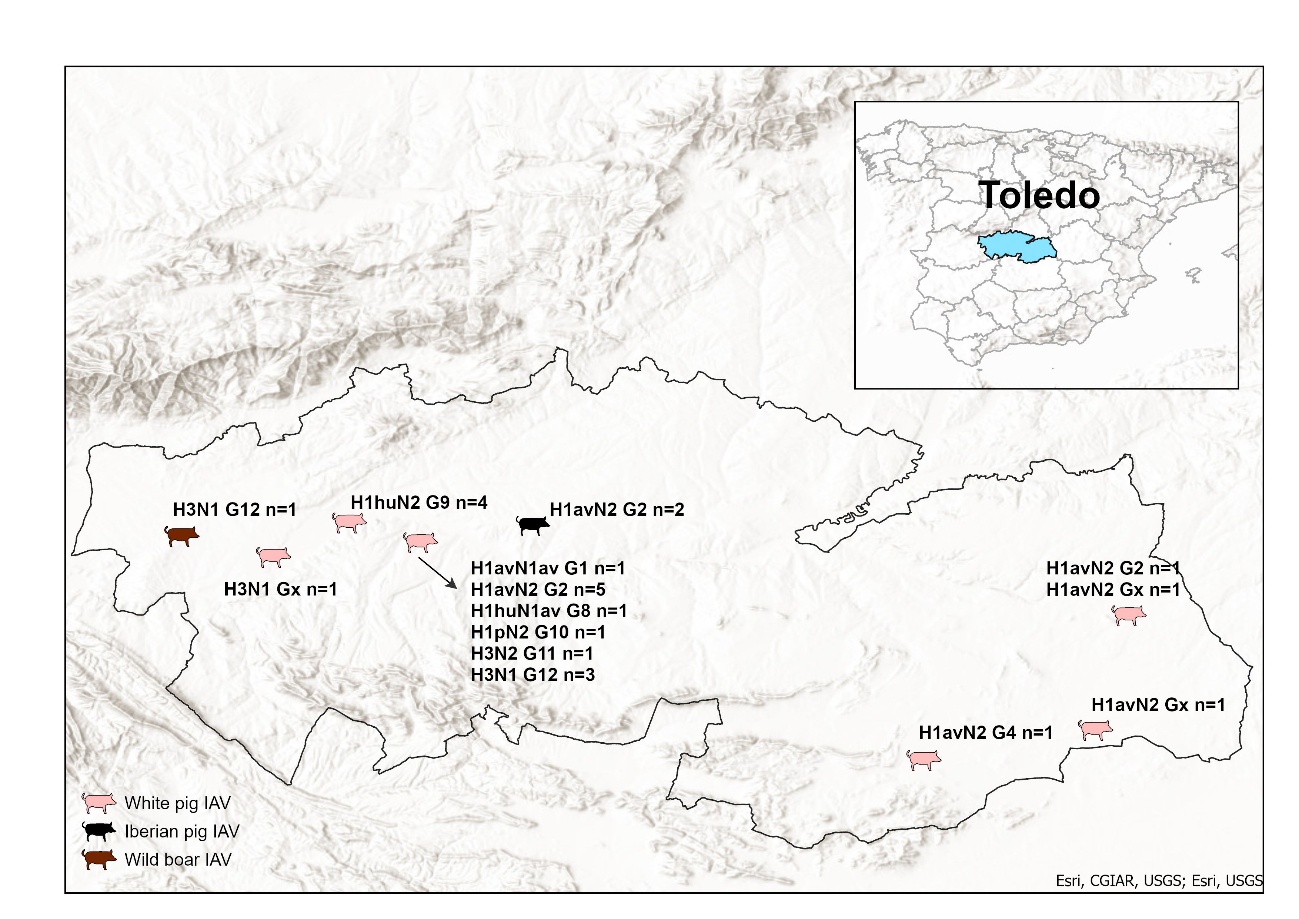

**Figure S2 Geographical distribution of the IAV strains detected in commercial Iberian (black) and white pigs (pink) and wild boar in the 2016-2022 period in Toledo.** G: genotypes detected within each subtype, av: Eurasian lineage, hu: human, p: pandemic, x: undetermined, n: number of isolates within genotype.

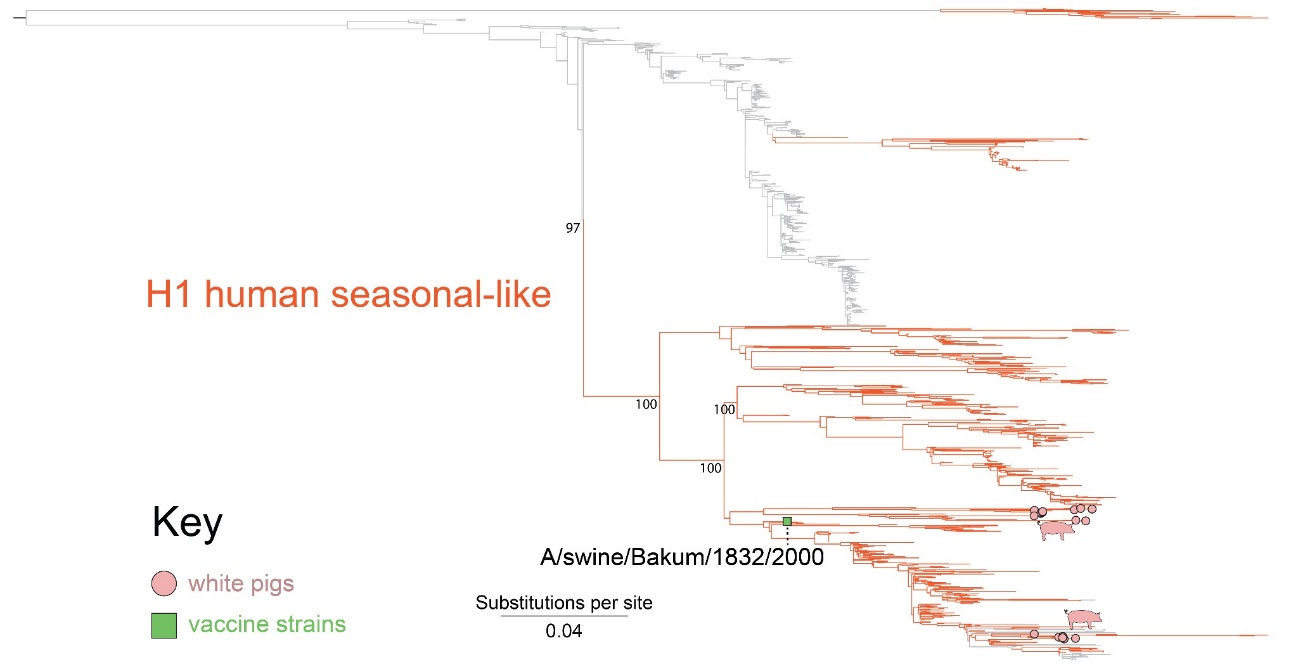

**Figure S3. Phylogenetic reconstruction of human and swine European H1 seasonal sequences from 1968-2008**. Gray branches represent viruses from humans in Europe while orange branches represent viruses from swine in Europe. Branch tips representing sequences obtained from commercial white pigs are annotated with pink circles. Vaccine strains are represented as green boxes with the vaccine strain name. Bootstrap values for lineages are included.

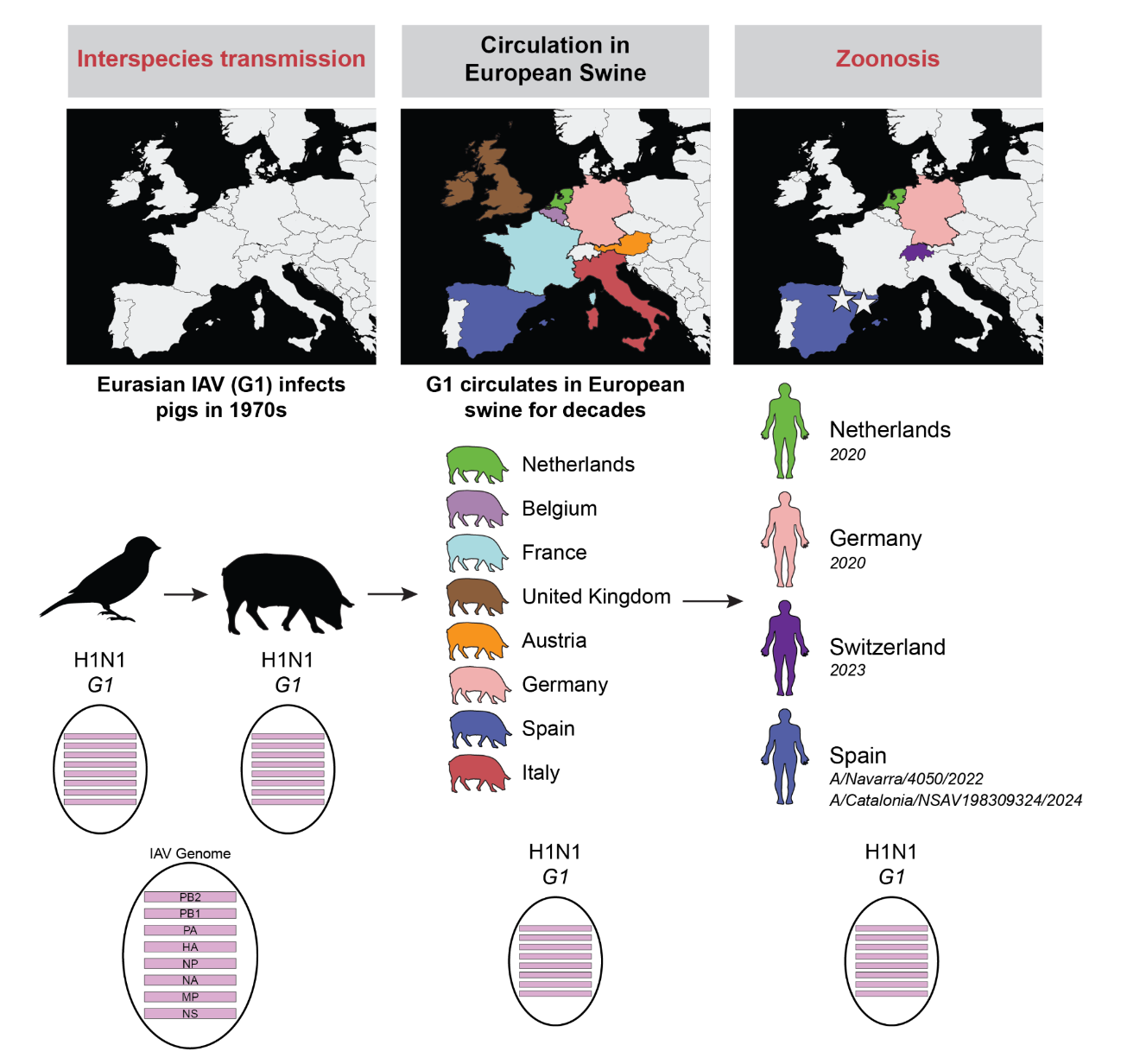

**Figure S4. Evolutionary history of the IAV genotype G1 associated with human zoonoses.** Genotype 1 (G1) originally transmitted from birds into European swine in the 1970s. G1 circulated in Europe for decades and was detected in multiple countries. Between 2020 and 2024, five variant cases were associated with G1 in Europe, two of them occurred in Spain A/Catalonia/NSAV600497186/2026 (H1N1v) and A/Catalonia/NSAV198309324/2024 (H1N1v).
